# Understanding the mechanisms underlying the lack of response to Janus kinase inhibition in ulcerative colitis

**DOI:** 10.1101/2024.10.17.618410

**Authors:** Elisa Melón-Ardanaz, Marisol Veny, Ana M. Corraliza, Alba Garrido-Trigo, Victoria Gudiño, Ángela Sanzo-Machuca, Marc Buendia, Miriam Esteller, Maite Rodrigo, M. Carme Masamunt, Ángel Giner, Ingrid Ordás, Agnès Fernández-Clotet, Berta Caballol, Ángel Corbí, Bram Verstockt, Severine Vermeire, Julian Panés, Elena Ricart, Azucena Salas

## Abstract

Ulcerative colitis (UC) is a chronic inflammatory disease of the colon. About one-third of UC patients failed to respond to available drugs, including tofacitinib, a broad Janus kinase (JAK) inhibitor. However, the mechanisms underlying patient response or resistance to oral JAK inhibitors remain unknown.

To elucidate the molecular and cellular pathways activated by tofacitinib in responder and non-responder patients, we generated a longitudinal single-cell RNA sequence dataset profiling both immune and non-immune cell populations from colonic biopsies of UC patients. Our analysis revealed that responders exhibited higher baseline JAK-STAT activity, while non-responders had increased baseline NF-kB pathway activation. Longitudinal comparisons showed that disease progression in non-responders was associated with increased abundance and enhanced activation of macrophages and fibroblasts. Our data suggest that resistance to tofacitinib is mediated by the hyperactivation of myeloid cells, and we identified IL-10-dependent macrophages as a cellular subset contributing to this resistance.

## INTRODUCTION

Ulcerative colitis (UC) is a chronic remitting and relapsing inflammatory bowel disease (IBD) that is characterized by superficial mucosal inflammation of the colon. Despite current access to biologics, including anti-TNF, anti-α4β7 and anti-IL-12/IL-23, about one third of patients continue to suffer from intestinal inflammation, which increases the risk of surgery (1) and long-term disability (2). In this context, more recently developed therapies, including oral inhibitors of the Janus kinases (JAK) family (3), show promise as they simultaneously target a broad spectrum of inflammatory signals that act on multiple cell types (4).

JAKs, which include JAK1, JAK2, JAK3 and TYK2, are a family of intracellular kinases that are essential to mediating responses to a large number of cytokines and growth factors (5). Tofacitinib, which blocks the catalytic functions of JAK1, JAK3 and, to a lesser extent, JAK2, was the first JAK inhibitor approved for the treatment of moderate-to-severe UC and psoriatic arthritis. Initially, tofacitinib was assumed to act primarily by blocking lymphocyte activation, based on its ability to block cytokines such as interleukin (IL)-2, IL-15, IL-4 or IL-7. However, it is now well accepted that tofacitinib (6) acts on most cell types beyond T and B lymphocytes, including non-hematopoietic (epithelial and stromal cells) and immune innate cell types (neutrophil, macrophages, dendritic cells, eosinophils, etc.). In fact, it inhibits a wider range of cytokines, including interferons (IFNs), IL-6, granulocyte-monocyte colony-stimulating factor (GM-CSF), and IL-10 family cytokines. Given its broad target profile, the effects of tofacitinib on different cell types in response to various stimulatory conditions may vary significantly, which could help explain the differences in efficacy seen in clinical practice, with almost 40% of UC patients not responding to treatment (3, 7). In this context, a better understanding of the mechanisms driving refractoriness is crucial to implement evidence-based therapeutic decisions and to design of alternative therapies. Here, we used single-cell RNA sequencing (scRNA-seq) analysis of intestinal cells from UC patients receiving tofacitinib to measure baseline and post-treatment differences in cell abundance, transcriptional states including inferred JAK-STAT signaling, and receptor-ligand interactions that could differentiate responder from non-responder patients.

Using this approach, we detected an overall enhanced baseline JAK-STAT pathway activity in responders, while increased baseline NF-KB activity was associated with non-responders. In addition, we identified LPS-activated IL-10-dependent macrophages as the main cellular subset potentially driving resistance to this treatment.

## RESULTS

### Response to tofacitinib in a real-life cohort of moderate-to-severe ulcerative colitis

Thirty-one patients with moderate-to-severe UC starting tofacitinib treatment as the standard-of-care participated in this observational study (Table 1; Figure 1A). Response to tofacitinib, evaluated based on endoscopic and clinical criteria as defined in the Methods sections, was achieved in 40% of patients (Table 1). Responders (R) showed a significant decrease in their histologic severity scores (Figure 1B) as defined in Supplementary Methods and in the number of plasma cells, lymphocytes, granulocytes (neutrophils and eosinophils), and stromal cells (endothelium and fibroblasts) as assessed in fixed tissues by an expert pathologist (MR) (Supplementary Figure 1). Remarkably, histological severity was significantly greater in the non-responders (NR), suggesting a worsening of the disease in this patient group (Figure 1B). Treatment with tofacitinib did not change the number of circulating leukocytes regardless of efficacy (Supplementary Figure 2A). In addition, we found no significant differences in the serum concentrations of tofacitinib when comparing R and NR (2.01 ± 2.38 *vs* 1.93 ± 2.45 ng/mL, respectively; p= 0.7717 Wilcoxon test). Activation of JAK signaling can be measured by detecting phosphorylation of their substrates (STATs). However, phosphorylation is transient and challenging to measure reliably in patient-derived samples. As a surrogate for JAK pathway activity in patient-derived samples, we measured the expression of JAK-STAT target genes (genes regulated downstream of JAK activation and STAT phosphorylation). Indeed, we found a significant inverse correlation between serum tofacitinib concentrations and the expression of the JAK-target genes *SOCS1*, *SOCS3* and *IRF1* in whole blood (Supplementary Figure 2B), suggesting target engagement. A similar trend was observed in biopsy samples, though this did not reach significance, potentially due to the smaller sample size (Supplementary Figure 2C).

**Figure 1.**
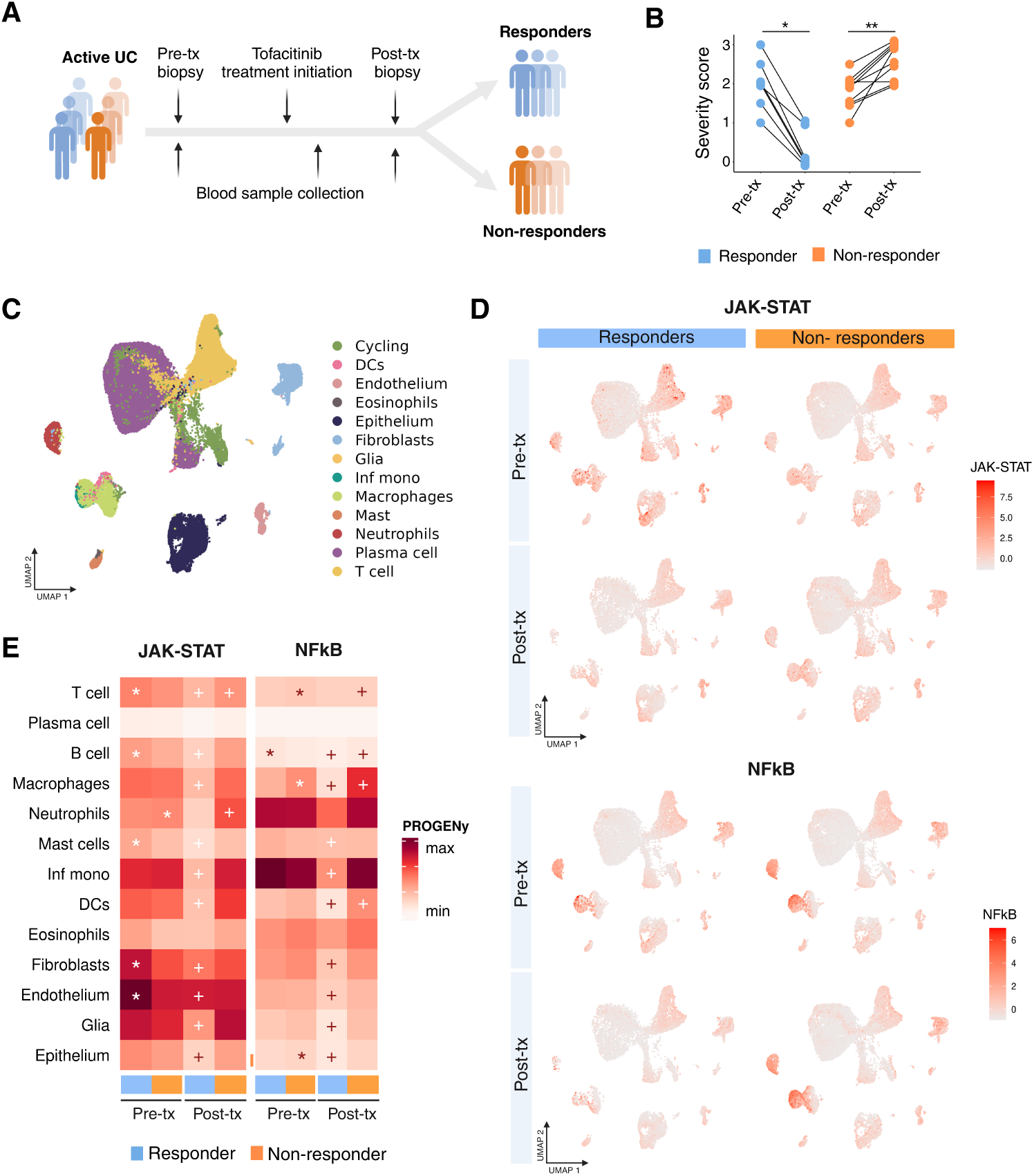
Study design and inferred pathway activity using scRNA-seq. **(A)** Schematic representation of the study workflow. Biopsies (for single-cell RNA-seq, transcriptional analysis and histological validation) and blood were collected from 31 patients before starting tofacitinib treatment (pre-tx) and during follow-up (post-tx). **(B)** Scores of histological severity in biopsies from responder and non-responder patients before and after tofacitinib treatment. Wilcoxon test for matched data (two-tailed p-value). *p<0.05, **p<0.01. **(C)** Uniform manifold approximation and projection (UMAP) of 69,813 intestinal cells color-coded by main cell type. **(D)** UMAPs showing the PROGENy pathway scores for JAK-STAT and NFkB separated by condition. **(E)** Heatmap showing the mean of JAK-STAT and NFkB pathway scores using PROGENy across all cell types in pre-and post-treatment samples. Wilcoxon signed-rank test adjusted using Bonferroni correction: *p<0.05 (R Pre-tx *vs* NR Pre-tx; * is shown for the group with increased higher mean pathway activation); + p<0.05 (Pre-tx vs Post-tx in R and NR).

**Table 1.** Clinical and demographic characteristics of study participants. Patient groups are separated by their response to tofacitinib.

|  | Responders |  | Non-Responders |  |
| --- | --- | --- | --- | --- |
| N | 14 |  | 17 |  |
| Sex (males/females) | 8/6 |  | 9/8 |  |
| Age (median, range, years) | 43 (22-67.6) |  | 48 (18.1-73.6) |  |
| Disease duration (median, range, years) | 8 (2-25.6) |  | 9 (1.9-28) |  |
| UC extension at baseline<br>(Pancolitis/Left-sided colitis/Proctitis) | 6/6/2 |  | 7/6/4 |  |
| Treatment at baseline |  |  |  |  |
| Aminosalicylates (e.g., mesalazina) | 5 |  | 4 |  |
| Corticosteroids (e.g., beclometasona) | 2 |  | 4 |  |
| Immunomodulator (e.g., Azathioprine) | 2 |  | 4 |  |
| Monotherapy (one of the above) | 7 |  | 8 |  |
| Combination therapy (two or more of the above) | 1 |  | 2 |  |
| Number of previous biologics (median, range) | 2 (0-4) |  | 2 (1-3) |  |
|  | Week 0 | Follow-up | Week 0 | Follow-up |
| CRP (mean, SD) | 0.48 (0.26) | 0.3* (0.2) | 0.93 (0.91) | 1.37 (1.34) |
| Endoscopic Mayo (median, range) | 3 (2-3) | 1 (0-3)** | 3 (2-3) | 3 (2-3) |
| Total Mayo (median, range) | 9 (4-12) | 3 (0-8)** | 10 (5-12) | 9 (6-12) |

\*p-value < 0.05; \*\*p-value < 0.01. N, number. CRP, C-reactive protein.

### Baseline JAK-STAT pathway activity is higher in patients that will favorably respond to treatment with tofacitinib

We generated scRNA-seq data (Figure 1C) from 23 colonic samples (taken at baseline and during follow-up) from a subset of patients (n=11) in the Barcelona cohort (Supplementary Table 1). Cells were annotated into 6 different clusters (Supplementary Figure 3A) and then further sub-clustered, resulting in 66 different cellular types/states (Supplementary Figure 3B). Markers used for the annotation of each cell subset are shown in Supplementary Methods.

The signaling activity of a particular pathway can be inferred using gene expression. Methods such as Pathway RespOnsive GENes (PROGENy) can accurately infer pathway activity, including JAK-STAT, from gene expression across a wide range of conditions (8). Importantly, this tool can be applied to scRNA-seq data, inferring pathway activity in a cell-by-cell manner. Using this approach, we measured the inferred activity of the JAK-STAT and NF-KB pathways on each intestinal cell cluster in R and NR patients at baseline, and after receiving tofacitinib treatment for at least 8 weeks (Figure 1D and E). Baseline JAK-STAT activity was significantly higher in R compared to NR across T cells, B cells, mast cells, fibroblasts and endothelial cells. In contrast, NR showed increased baseline NF-KB pathway activity in T cells, macrophages and epithelial cells. Importantly in R patients, both JAK-STAT and NF-KB pathway activity was significantly reduced following treatment, both in immune and non-immune cell types. In contrast, we observed a significant increase in NF-KB pathway activity (438.17 w0 vs 1016.07 post-tx, q-value 3.76×10^-47^) in macrophages from NR patients who had received tofacitinib.

### Treatment with tofacitinib induces significant changes in the cellular composition of the intestinal mucosa

In addition to changes in JAK-STAT and NF-kB pathway activity, response to tofacitinib was associated with significant changes in the abundances of different cell types, including decreases in myeloid, B and plasma cells, and increases in epithelial and stromal populations (Figure 2A and 2B). In contrast, NR patients treated with tofacitinib did not experience epithelial or stromal recovery. Nor did tofacitinib reduce the abundance of myeloid, plasma cells, B cells, cycling cells or inflammatory fibroblasts. Nonetheless, lack of response to tofacitinib was unexpectedly associated with a significant increase in the abundance of myeloid cells. Analysis of the specific changes within this compartment revealed that neutrophils, inflammatory INHBA+ macrophages, heat shock macrophages, inflammatory monocytes and dendritic cells were all significantly expanded in NR following treatment with tofacitinib (Figure 2B).

**Figure 2.**
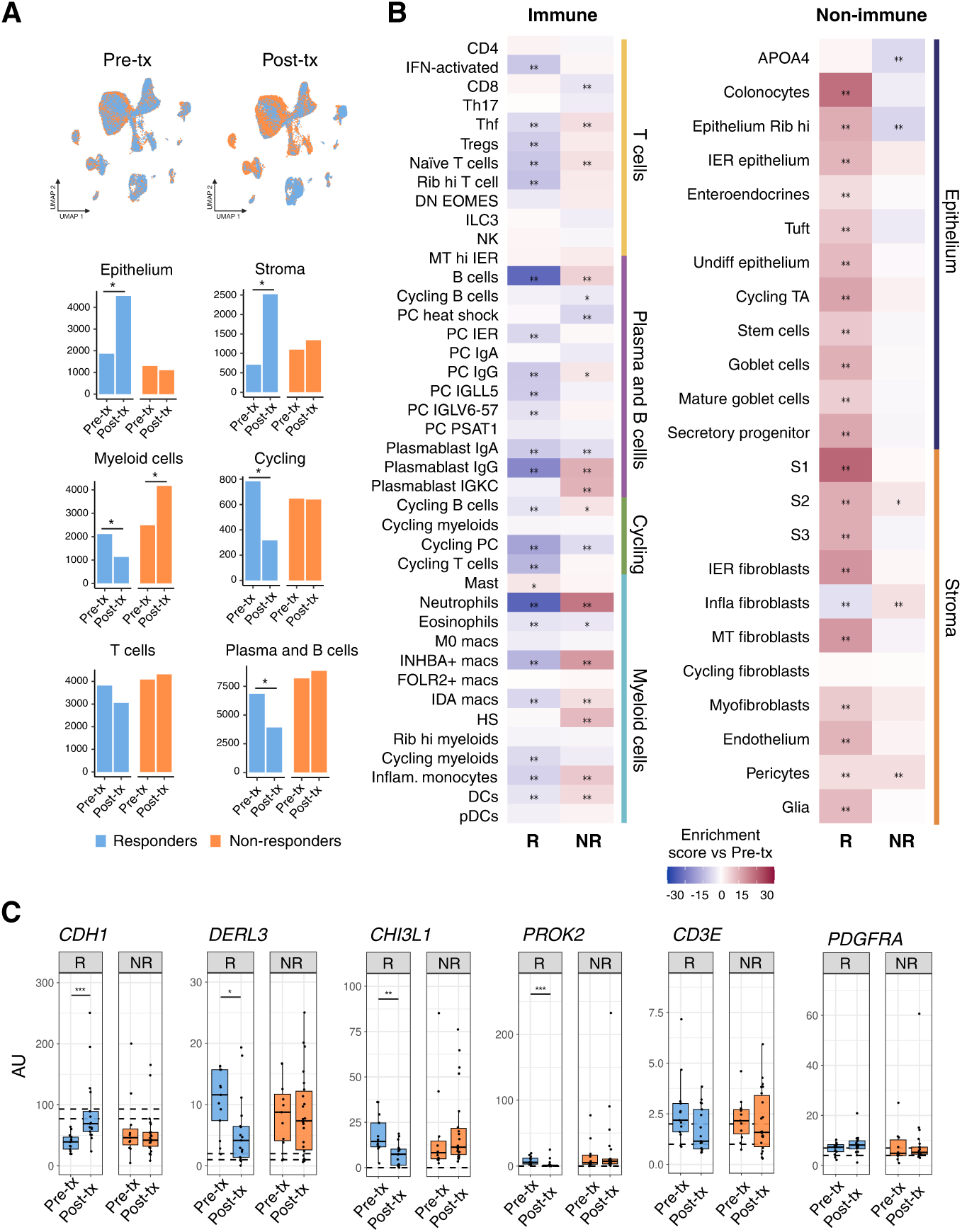
Treatment with tofacitinib induces significant changes in the abundance of intestinal cell types in responder and non-responder patients. **(A)** UMAPs of pre-treatment (pre-tx; 33,941 cells) and post-treatment (post-tx; 35,872 cells) intestinal cells color-coded by response to tofacitinib. Bar plots depict the cellular abundance of the six major cell types (epithelium, stroma, plasma and B cells, myeloid cells, cycling and T cells) for responder and non-responder patients before (pre-tx) starting tofacitinib and during follow up (post-tx). Comparison was performed using the Chi square test *p<0.05. **(B)** Intestinal cell abundance is shown as the enrichment scores of immune (T cells, plasma and B cells, cycling and myeloid cells) and non-immune (epithelium and stroma) sub-clusters in responders and non-responders relative to baseline (pre-tx). Comparisons were performed using the Chi square test *p<0.05 and **p<0.01. **(C)** Gene expression analysis in whole-biopsy bulk RNA. Selected genes were markers for the main cell lineages. Data is shown separately for responders (blue) pre-tx (n=13 samples) and post-tx (n=17 samples); and for non-responders (orange) pre-tx (n=11 samples) and post-tx (n=22 samples). Data is expressed as arbitrary units (AU) and presented as boxplots. Dotted lines show the standard error of the mean of biopsies from healthy controls (n=10). Wilcoxon test. False discovery rate-corrected p-values: *< 0.05; **< 0.01 and ***< 0.001.

Next, we measured the expression of selected cell-type and cell-state gene markers in intestinal biopsies from the Barcelona cohort (n=14 patients, 33 samples), as well as additional samples collected at a different center (Leuven cohort, n=10 patients, 30 samples) (Supplementary Table 1). Analysis of gene expression changes using bulk RNA from biopsies, while lacking single-cell resolution, did detect changes in the overall proportions of cell types that display unique gene markers. Indeed, changes in the mRNA expression of epithelial (*CDH1* and *TFF3*), plasma cell (*DERL3*), inflammatory fibroblasts (*CHI3L1*), neutrophils (*PROK2*), monocytes (*FCGR3A*) and inflammatory markers (*CXCL10* and *S100A8*) mirrored changes in cell proportions, as revealed by scRNA-seq (Figure 2C and Supplementary Figure 4). Also, in agreement with the scRNAseq findings, no change in the T-cell marker *CD3E* was detected using bulk mRNA. In contrast, bulk RNA analysis could not capture the changes observed by scRNA-seq in NR patients, such as the expansion of the myeloid compartment following tofacitinib treatment (Figure 2C and Supplementary Figure 4). In summary, we conclude that scRNA-seq can reveal changes in cell-specific pathway activity, cell proportions and cell states, some of which are not readily detectable by bulk analysis, thus providing a clear advantage when trying to disentangle changes in cellular states between patient groups.

### Tofacitinib treatment shapes the transcriptional signatures of intestinal macrophages

In addition to changes in the proportions of a particular cell population (Figure 3A), scRNA-seq can detect variations in gene expression within a given cell type when comparing groups of samples. Following tofacitinib treatment, differentially expressed genes (DEG) were measured in both R and NR for all macrophage populations (Figure 3B, Supplementary Figure 5A and Supplementary Table 2A). FOLR2+ macrophages showed the highest number of regulated genes, with 533 of them significantly regulated (95% upregulated; FC>1.5, FDR>0.05) following the response to tofacitinib (Figure 3B, Supplementary Table 2A). Upregulated genes included key metabolic and immune tolerogenic genes such as *IGF1*, *CD163L1*, *CLEC10A*, *MAF*, *AHR* and *IL10RA*. The most significantly downregulated gene was *S100A9*, which encodes for one of the calprotectin subunits. Other genes such as *FCGR3A*, *CCL13*, *MMP12*, *FCGR2A* and IFN-response genes such as *STAT1*, *IFITM3*, *GBP1* and MHC Class I were also significantly downregulated in R. Changes in M0 and IDA macrophages were less pronounced but included the significant downregulation of *S100A9* and the IFN-response genes *STAT1*, *ISG15*, *IFI27*, MHC Class I genes and *GBP1* (Supplementary Figure 5A and Supplementary Table 2A). Overall, these results demonstrated an attenuation in the inflammatory signatures of macrophages, together with the recovery of a healthy homeostatic intestinal resident macrophage signature in R.

**Figure 3.**
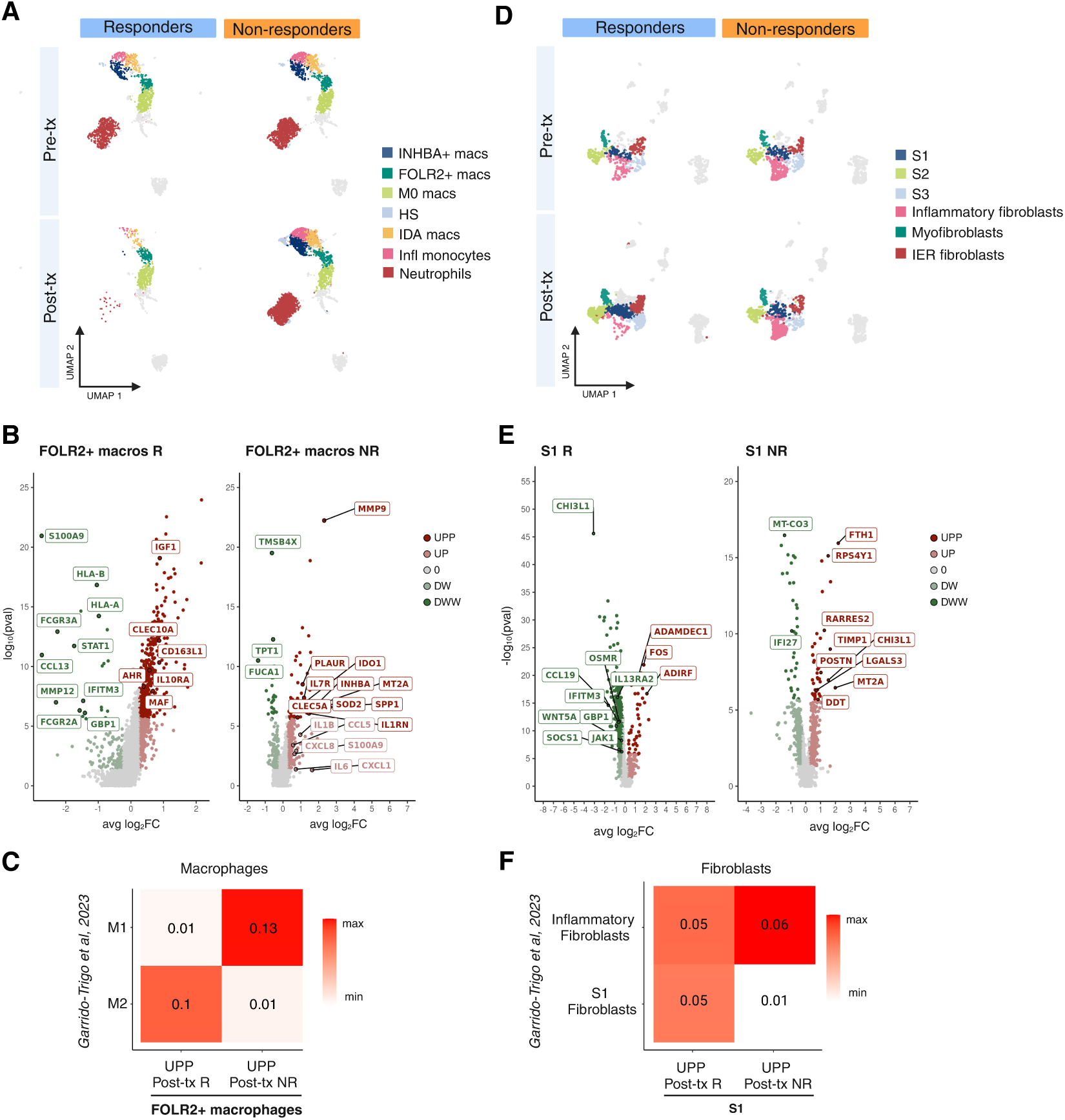
Tofacitinib induces significant transcriptional changes in intestinal macrophages and fibroblasts. **(A)** Uniform manifold approximation and projection (UMAP) of scRNA-seq data for myeloid cells colored by cell subsets (9,902 cells) from responders and non-responders before (pre-tx) and after (post-tx) treatment with tofacitinib. **(B)** Volcano plots showing genes differentially expressed in FOLR2+ macrophages post-tx compared to pre-tx in responder (left panel) and non-responder (right panel) patients. A two-sided Wilcoxon rank sum test was applied. Genes with a false discovery rate adjusted p value < 0.05, and a fold change (FC) > 1.2 (UPP, dark red) or FC < 0.83 (DWW, dark green) were considered regulated. Additionally, genes with a nominal p-value < 0.05 are shown in light red (UP) if FC > 1.2 and light green if FC < 0.83. macros, macrophages. **(C)** Jaccard analyses comparing significantly upregulated (adjusted p.value < 0.05 & FC > 1.2, UPP) genes in FOLR2+ macrophages post-tx in responder and non-responder patients with canonical markers of M1 or M2 macrophages from Garrido-Trigo *et al* (43). **(D)** UMAP of scRNA-seq data for stromal cells colored by cell subsets (5,674 cells) from responders and non-responders before (pre-tx) and after (post-tx) treatment with tofacitinib. **(E)** Volcano plots showing genes differentially expressed in S1 fibroblasts post-tx compared to pre-tx in responders (left panel) and non-responders (right panel). A two-sided Wilcoxon rank sum test was applied. Genes with a false discovery rate adjusted p value < 0.05, and a fold change (FC) > 1.2 (UPP, dark red) or FC < 0.83 (DWW, dark green) were considered regulated. Additionally, genes with a nominal p-value < 0.05 are shown in red (UP) if FC > 1.2 and green if FC < 0.83. **(F)** Jaccard analyses comparing significantly upregulated (adjusted p.value < 0.05 & FC > 1.2, UPP) genes in S1 fibroblasts post-tx in responder and non-responder patients with canonical markers of S1 or inflammatory fibroblasts from the study by Garrido-Trigo *et al* (43).

Compared to R, the number of genes significantly regulated by tofacitinib treatment in macrophages was lower in the NR population (Figure 3B, Supplementary Figure 5B and Supplementary Table 2B). Again, FOLR2+ macrophages showed the highest number of DEG (84 genes, 56 upregulated) after tofacitinib treatment. Overall, in NR tofacitinib significantly increased the expression of genes normally associated with an inflammatory phenotype, including *MT2A*, *IDO1*, *SPP1*, *CLEC5A*, *MMP9*, *PLAUR*, *IL1RN*, *IL7R* and *SOD2* (Figure 3B).

Indeed, FOLR2+ macrophages in R and NR patients showed opposing patterns following treatment with tofacitinib, with macrophages in refractory patients acquiring inflammatory features (Figure 3C). Taken together, these results show that tofacitinib significantly regulates the transcriptional signatures of macrophages, with opposing effects in R and NR patients.

### Response to tofacitinib is associated with significant remodeling of the fibroblast compartment

Similar to macrophages, tofacitinib also induced significant changes in the composition and transcriptional patterns of fibroblasts (Figure 2B, Figure 3D and Supplementary Figure 6). In R, S1 (lamina propria *ADAMDEC*) fibroblasts showed the largest number of DEG (456 genes, 416 downregulated) in response to tofacitinib (Supplementary Table 3A). Downregulated genes in R included *CHI3L1*, *HIF1A*, *WNT5A*, IFN-response genes *IFI27*, *IFITM3*, *ISG15*, *GBP1*, *CCL19*, the IFN signaling components *JAK1* and *STAT1*, and its suppressor *SOCS1*, as well as the JAK-dependent cytokine receptor chains *IL7R*, *IL4R*, *OSMR*, *IL15RA* and *IL13RA2* (Figure 3E). Analysis of the most significantly regulated pathways in S1 fibroblasts from R emphasizes the ability of tofacitinib to regulate key immune and tissue remodeling pathways (Supplementary Figure 7). Moreover, in R the abundance of inflammatory fibroblasts was significantly lower after treatment with tofacitinib (Figure 2B and Figure 3D).

Compared to R, in NR the number of DEG in fibroblasts following tofacitinib treatment was much lower (Supplementary Table 3B). In agreement to what we observed in FOLR2+ macrophages, S1 fibroblasts showed a heightened inflammatory profile following tofacitinib treatment. Upregulated genes included a few associated with inflammatory states, such as *CHI3L1*, *TIMP1*, *POSTN*, and *MT2A* (Figure 3E and 3F). Hence, following tofacitinib treatment S1 fibroblasts showed opposing gene regulation profiles in R and NR, though to a lesser degree than what we observed in macrophages.

### Tofacitinib increases the expression of inflammatory genes in lipopolysaccharide-stimulated macrophages

To explain the discrepancy in the impact of tofacitinib on macrophage and fibroblast activation when comparing R and NR patients, we explored the potentially different effects of tofacitinib in cell cultures exposed to diverse stimuli. As shown earlier, macrophages in NR showed increased inferred NF-kB pathway activity (Figure 1D and E), which could be driven by stimuli such as TNF or LPS, among others, whereas the increased JAK-STAT signaling detected in fibroblasts from responders matched that of IFNγ, or other JAK-dependent signals. We therefore generated cultures of monocyte-derived macrophages (Figure 4A), or primary intestinal fibroblasts (Figure 4B) treated with NF-kB-dependent (bacterial LPS or TNF) or JAK-dependent (IFNγ) inflammatory mediators. Transcriptional changes induced by each stimulus (mean log2fold_change compared to unstimulated conditions) (Figures 4C and 4D, Supplementary Figure 8) and the effects of tofacitinib at concentrations similar to the Cmax detected in serum (Figure 4E and 4F) are shown for both macrophages and fibroblasts. Tofacitinib had an overall inhibitory effect on the response of macrophages to TNF or IFNγ. In contrast, in the context of LPS stimulation of macrophages, tofacitinib significantly upregulated the transcription of *IDO1*, *CXCL1*, *CXCL8, CCL5*, *IL1B*, *IL6, IL23*, *INHBA, CLEC5A* and *MMP9,* among other genes, compared to LPS only stimulated macrophages. The increases in TNF, CXCL1 and IL-6 protein secretion in culture supernatants from LPS/tofacitinib-treated macrophages were confirmed by ELISA (Figure 4G). Tofacitinib did, however, significantly downregulate the expression of *SOCS3* and *ACOD1*, as well as the secretion of CXCL10 in LPS-stimulated macrophages, confirming adequate target engagement.

**Figure 4.**
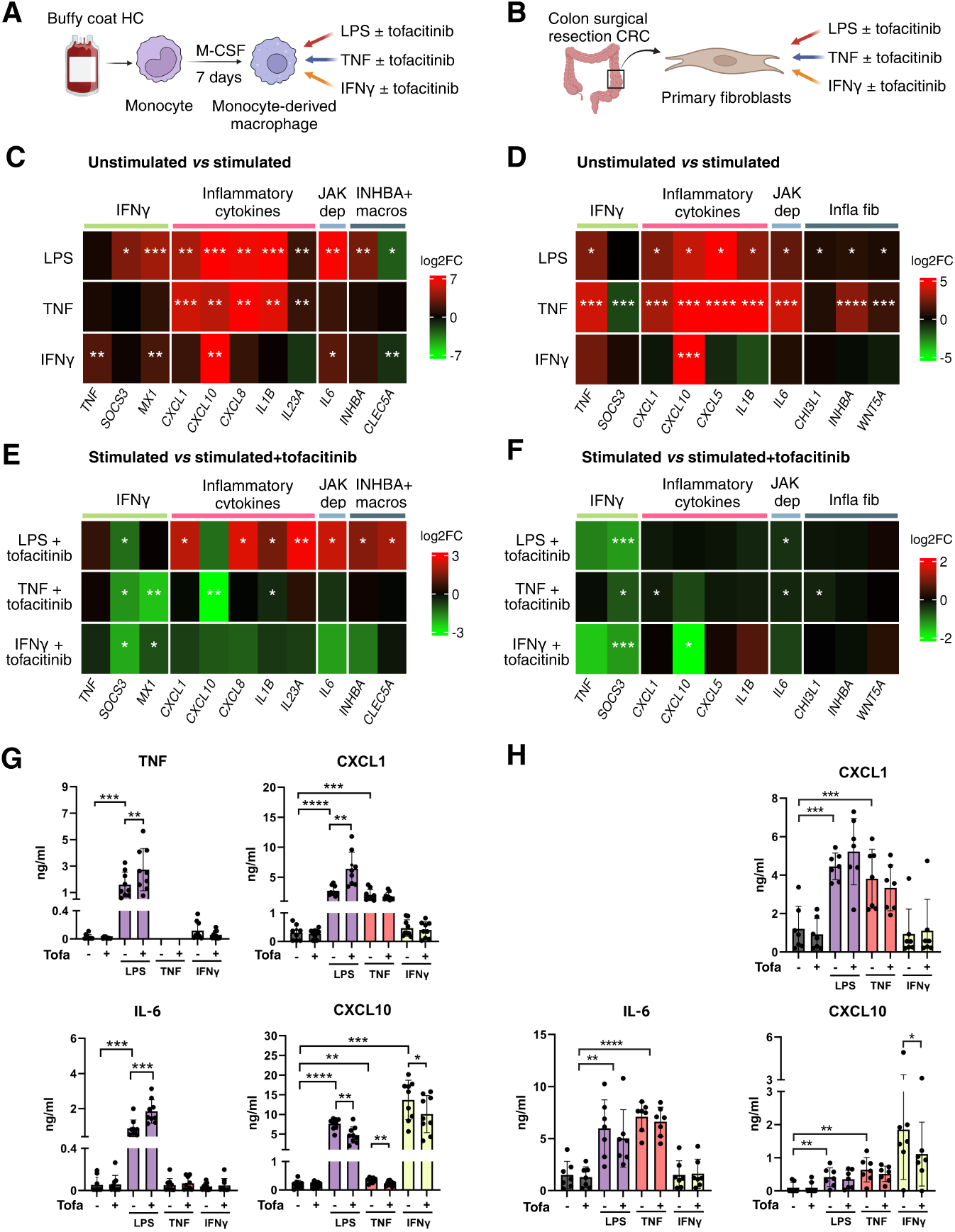
Differential effects of tofacitinib on *in vitro-*activated macrophages and fibroblasts. Experimental design: M-CSF-monocyte-derived macrophages **(A)** or intestinal fibroblasts **(B)** were exposed to different inflammatory stimuli (LPS, TNF or IFNγ) in the presence/absence of tofacitinib. Total RNA and supernatants were collected after 19 h of culture. Heatmaps showing mean log2 of fold-change (log2FC) in the expression of selected genes. Changes in response to LPS (10 ng/ml), TNF (20 ng/ml) and IFNγ (5 ng/ml) compared to an unstimulated control are shown for **(C)** M-CSF-monocyte-derived macrophages and **(D)** fibroblasts. Changes in gene expression induced by tofacitinib (300 nM) on LPS, TNF or IFNγ−stimulated M-CSF-monocyte-derived macrophages **(E)** or fibroblasts **(F)**. n=5 and n=6 independent experiments, respectively. One-sample t-test false discovery rate-corrected p-values: *<0.05, **<0.01, ***<0.001, ****<0.0001. Concentrations of cytokines in supernatants of M-CSF-monocyte-derived macrophages **(G)** or fibroblasts **(H)** exposed to the different inflammatory stimuli (LPS, IFNγ or TNF) in the presence of tofacitinib or DMSO as vehicle control. n=9 and n=7 independent experiments, respectively. Data are expressed as median±range. Paired t-test false discovery rate-corrected p-values: *<0.05, **<0.01, ***<0.001, ****<0.0001. JAK dep, JAK dependent. INHBA+ macros, INHBA+ macrophages. Infla fib, inflammatory fibroblasts.

In contrast to macrophages, fibroblasts showed an overall down regulation of gene expression in response to tofacitinib regardless of the stimuli (Figure 4F and 4H). Nonetheless, when fibroblasts were exposed to supernatants from LPS/tofacitinib-treated macrophages, they upregulated the expression of the pro-inflammatory mediators *CXCL1*, *CXCL5*, *IL1B* and *IL6* (Supplementary Figure 9A and B).

In conclusion, *in vitro* experiments confirm that tofacitinib can induce markedly different states on macrophages depending on the stimulatory mediator. This effect was not detected in fibroblasts under the same stimulatory conditions. However, we propose that fibroblasts can sense changes in macrophage activation, which may result in their heightened response *in vivo*.

### Tofacitinib may enhance responses to LPS by interfering with IL-10 signaling on macrophages

In response to LPS, monocyte-derived macrophages generated in the presence of M-CSF secrete abundant IL-10 (Figure 5A) (9, 10), which regulates the overall pro-inflammatory response to bacterial sensing. Interfering with the autocrine effects of IL-10 in the context of LPS stimulation results in increased macrophage (11) and monocyte (12) activation. In contrast, primary fibroblasts did not produce detectable levels of IL-10 in response to LPS (data not shown). Given that IL-10 signaling is a JAK-dependent pathway, we hypothesized that hyperactivation of LPS-stimulated macrophages, when treated with tofacitinib, would be mediated by the inhibition of IL-10 signals. Indeed, several of the genes that we found to be enhanced by tofacitinib on LPS-treated macrophages were previously described to be regulated by IL-10 in LPS-stimulated macrophages (13). This suggests that the paradoxical effects of tofacitinib on LPS-stimulated macrophages may rely on the inhibitor’s ability to block signaling downstream of IL-10. To determine whether the heightened macrophage activation in the mucosa of NR may also be the result of IL-10 signaling inhibition, we took advantage of a published RNA-seq dataset containing M-CSF-derived macrophages stimulated with LPS in the presence of an anti-IL-10 antibody (11). Of the more than 200 genes that were significantly upregulated by IL-10 blockade, 24 were similarly regulated by tofacitinib in the intestinal FOLR2+ macrophages from NR patients (Pearson correlation, p=2.344×10^-5^) (Figure 5B).

**Figure 5.**
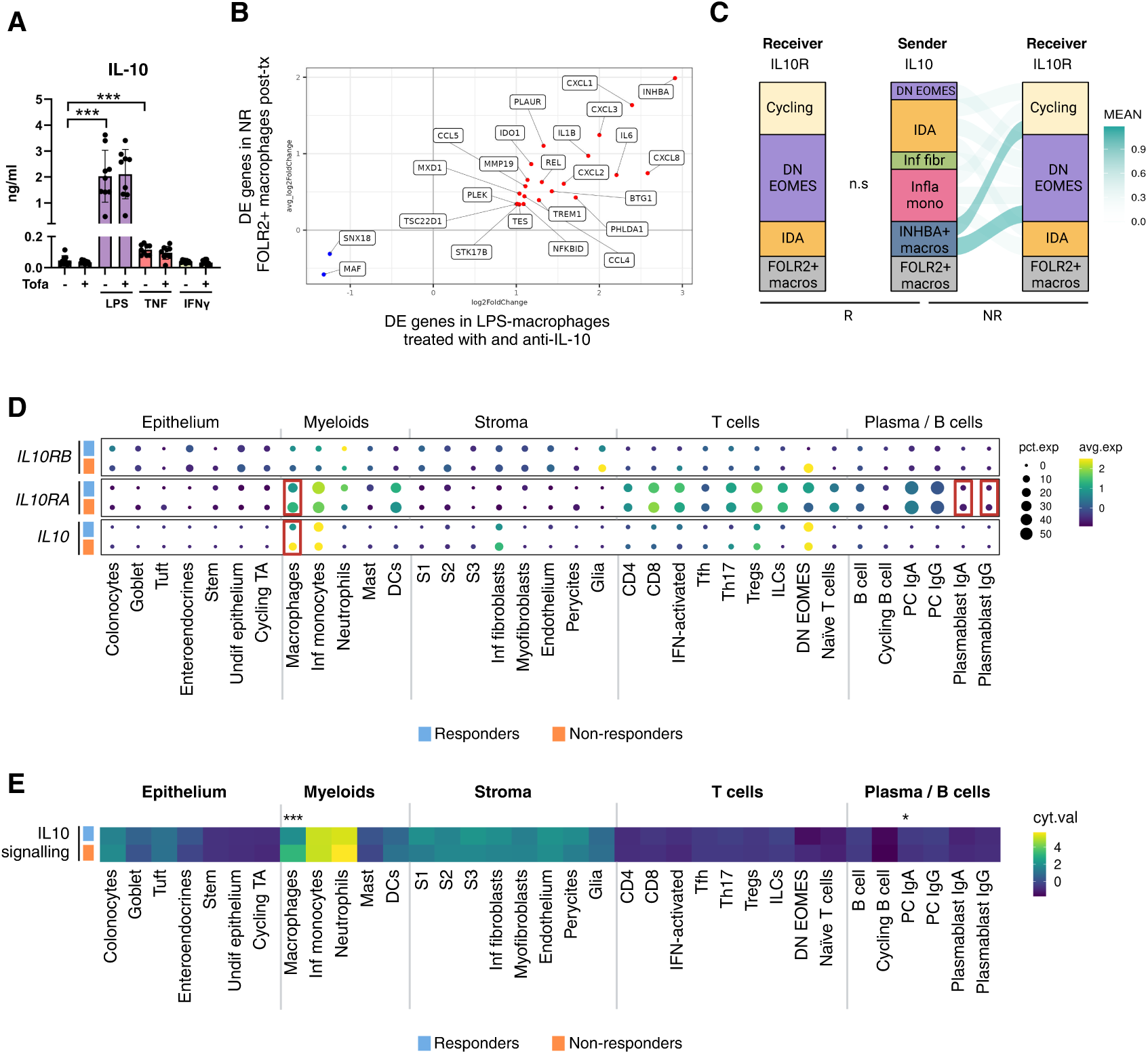
Tofacitinib may enhance responses to LPS by interfering with IL-10 signaling on macrophages. **(A)** Concentrations of IL-10 in supernatants of culture monocyte-derived macrophages under different stimulatory conditions. Macrophages were treated with vehicle (DMSO) or tofacitinib (300 nM). Data are expressed as median±range. Paired t-test false discovery rate-corrected ***p-value< 0.001. **(B)** Scatter plot of log2FC of genes regulated by anti-IL-10 mAb in LPS-stimulated macrophages (data from Cuevas *et al* (11) compared to the log2FC of genes regulated in FOLR2+ macrophages (scRNAseq data) from patients non-responsive to tofacitinib. Pearson test, p=2.344×10^-5^. **(C)** CellPhoneDB cell communication analysis using baseline data in responder (R) and non-responder (NR) patients. Results are presented as a galluvial plot, where lines connect communicating populations, and colors represent the mean intensity of the interactions. ns, not significant; macros, macrophages. **(D)** Mean expression of mRNA transcripts for each cell type is shown for *IL10RB*, *IL10RA* and *IL10* in pre-treatment inflamed samples from R and NR patients. Comparison was performed using Wilcoxon signed-rank test and adjusted using a Bonferroni correction. Significant differences (p<0.05) between responders and non-responders are shown in red. **(E)** IL-10 signaling scores (CytoSig) are shown for pre-treatment inflamed samples. Heatmap shows relative enrichment of IL-10 signaling scores in R and NR patients. Wilcoxon signed-rank test adjusted using Bonferroni correction was applied to compare R versus NR. * p<0.05 ** p<0.01, *** p<0.001.

Given the potential link between IL-10 blockade and worsening of disease after tofacitinib, we asked whether increased IL-10-IL-10R interactions were present at baseline (before starting tofacitinib treatment) in NR compared to R. CellPhoneDB (14) analysis (a tool that measures ligand-receptor interactions based on scRNA-seq expression data) detected significant interactions between IL-10-IL10R at baseline (week 0) in samples from NR, but not in R patients (Figure 5C). In addition, we measured the mean expression of *IL10* and its receptor chains *IL10RA* and *IL10RB* at baseline for R and NR (Figure 5D). IL-10 is mostly produced by myeloid cells, including macrophages, which showed significantly higher IL10 and IL10RA expression in the NR population. Finally, we applied CytoSig (15) to scRNA-seq expression data to measure the signaling activity of IL-10 in R and NR at week 0 (Figure 5E). This analysis confirmed significantly increased IL-10 signaling in macrophages from NR patients compared to R before starting tofacitinib. Overall, these results favor the hypothesis that IL-10 signaling is more prevalent in NR before starting tofacitinib treatment.

In conclusion, we show that in addition to increased NF-KB inferred signaling activity, increased *IL10* transcription, IL-10-IL-10R interactions and IL-10 signaling within macrophages prior to the start of tofacitinib treatment is associated with a lack of response to tofacitinib.

## DISCUSION

Lack of response to therapy remains an unpredictable event affecting about 30% of patients suffering from IBD, including UC (16). Despite extensive research, little is known about the potential predictors of drug failure or the mechanisms driving the lack of response in refractory patients. To predict or to understand refractoriness, most studies to date have focused on bulk transcriptomic analysis of baseline intestinal biopsies. While bulk RNA analysis has been instrumental in uncovering the molecular patterns mediating chronic intestinal inflammation (17–20), this technology cannot robustly differentiate active disease signatures from patients with different behaviors, including drug response. Although some baseline transcriptional signatures have been associated with refractory behaviors (18, 21–25), these have proven difficult to validate (26) and, as a result, none have yet been translated into clinical practice. More recently, scRNA-seq provides a high-resolution picture of cell identities (27–35) in a manner that bulk transcriptomics does not. Importantly, scRNA-seq can detect changes in gene expression within cell types across patient groups or time points, as supported by our data. For the first time, the single-cell transcriptomes of intestinal cells have now been measured in order to dissect the relevant cellular subsets and genes involved in response and/or resistance to tofacitinib, while also offering novel insights into the prediction of treatment response.

Analysis of the changes in gene expression within subsets of macrophages or intestinal fibroblasts provided important clues on how specific cell types differentially respond to JAK inhibition. For instance, we could measure how resident FOLR2+ macrophages and lamina propria S1 fibroblasts adopted opposite signatures in R and NR patients following treatment with tofacitinib. Furthermore, we applied computational tools to infer pathway activity, cell-specific signaling responses and receptor-ligand interactions at a single-cell resolution. None of these analyses would have been possible without applying a single-cell approach.

The paradoxical effects that tofacitinib treatment has on intestinal cells drove us to hypothesize that not all cell states may respond equally to JAK inhibition. Given the complexity and multiple factors that can result in JAK activation, tofacitinib would be expected to act on numerous pathways, some of them with divergent effects. Indeed, this family of kinases control the response to both regulatory and immune-activating cytokines (5). Importantly, and as shown here, JAK signaling is not only induced by direct activators such as IFNs, IL-6, IL-23, or IL-10, but also indirectly downstream of other key proinflammatory mediators such as TNF or toll-like receptor ligands such as LPS. While tofacitinib cannot directly interfere with signaling through TNF or LPS receptors, response to these mediators involves the production of several JAK-dependent cytokines including IL-6, IL-23, IL-12, or IL-10, which carry out most of the downstream effects. Overall, we hypothesized that this complex network of interactions, and the fact that both pro- and anti-inflammatory signals can be blocked by JAK-inhibition, could explain the variable efficacy observed in UC patients treated with tofacitinib.

Overall, our data emphasizes the key role that macrophages play in orchestrating responses to tofacitinib, and strongly suggests that IL-10 blockade of these cells in the context of ongoing inflammation may explain disease exacerbation in those patients who become refractory. Macrophages are, indeed, a well-known source of this cytokine in response to pathogen-associated molecular pattern signaling, and recent work has shown the key role played by IL-10 autocrine signaling in controlling the resolution to inflammatory responses in macrophages (13). In addition, while most hematopoietic cells (including lymphocytes) can express the IL-10 receptor, macrophages are critically regulated by their responses to this cytokine. In fact, IL-10 controls metabolic reprogramming, autophagy, mitochondrial dysfunction, and activation of the inflammasome on macrophages exposed to bacterial components (36). Specifically, in the context of the gut, lack of IL-10 signaling on macrophages can, in fact, drive severe intestinal inflammation (37, 38). In addition, IL-10 signaling on macrophages, but not T cells, has been found to be essential for a full therapeutic response to anti-TNF (39). Moreover, direct blockade of IL-10 exacerbates macrophage activation in response to microbial patterns (11, 13). All these data fit with our findings that the blockade of IL-10 signaling on LPS-stimulated macrophages could exacerbate their inflammatory potential.

Besides macrophages, we also observed increased activation of fibroblasts in NR patients following treatment with tofacitinib. However, fibroblast activation was not tightly controlled by IL-10 and was overall mediated by tofacitinib under the conditions used. Therefore, we suggest that the exacerbated activation of fibroblasts seen in NR could well be secondary to the lack of IL-10 control on resident macrophages. In particular, the increased production by macrophages of cytokines, such as IL-1β, TNF or IL-6, which are all potent activators of fibroblasts (25), would heighten the overall tissue inflammation.

A dual effect of tofacitinib has been reported in murine Th17 cells, depending on the conditions under which they were generated (40), mirroring what we report here in macrophages. When Th17 cells were differentiated in response to IL-6, IL-1β and IL-23, tofacitinib reduced their inflammatory potential. In contrast, when differentiated with IL-6 and TGFβ, conditions that drive the differentiation of IL-10 producing Th17 cells, treatment with tofacitinib increased their ability to secrete IL-17. This demonstrates that, under stimulating conditions in which IL-10 is produced and acts as a regulatory feedback loop, JAK inhibition leads to increased activation rather than immunosuppression.

To our knowledge, few studies have used single-cell transcriptomics to monitor the effects of drug response in patients (41). In one recent study, skin samples from a small cohort of patients with systemic sclerosis receiving tofacitinib were analyzed using this approach (42). While this study did not stratify samples based on response to treatment, it reported results similar to our own, namely that tofacitinib had significant effects on the stromal and myeloid compartments, with limited changes to the T- and B-cell subsets.

Our study has some limitations, including the relatively small sample size and the diverse follow-up times at which samples could be collected. In addition, while this study did not aim to identify predictors of drug response at baseline, we strongly believe that understanding the mechanisms of "drug escape" may be a more relevant objective, not only to ensure efficacy and limit drug failure, but also to potentially guide follow-up therapeutic decisions. Indeed, refractoriness to JAK inhibition led to increased numbers of neutrophils, inflammatory monocytes, and macrophages, as well as to the production of IL-6, TNF and CXCL1, among others, which suggests these could become potential therapeutic targets for those patients who fail to respond to a JAK inhibitor.

Overall, we demonstrate that tofacitinib can not only dampen macrophage and fibroblast activation *in vivo* and *in vitro*, but also help enable epithelial regeneration and reduce the numbers of plasma cells, certain subsets of T cells and the overall restitution of mucosal homeostasis in patients who respond to therapy. Thus, our data strongly validates the relevance of JAK inhibition in treating moderate-to-severe intestinal inflammation. However, our study also shows that resistance to tofacitinib is a remarkably active process that we propose is, at least in part, caused by the direct inhibition of IL-10 signaling in subsets of activated macrophages. While we cannot exclude the possibility that other mechanisms might play a role in refractory disease, including the effects of IL-10 inhibition on CD4 T cells, such as Th17 cells, here we provide the first evidence of a specific molecular pathway involved in the escape from JAK-driven immunosuppression.

## METHODS

### Sex as a biological variable

Sex was not considered as a biological variable; therefore, patients of both sexes were included in the study.

### Patient recruitment and treatment

Patients who were candidates to receive tofacitinib (10 mg BID) as a standard-of-care treatment in our IBD Unit (Barcelona cohort, Hospital Clinic, Barcelona, Spain) or UZ Leuven Hospital (Leuven cohort, Leuven, Belgium) were invited to participate in the observational study (Supplementary Table 1). Patients showing no endoscopic and/or clinical improvement at week 8 were maintained on tofacitinib 10 mg BID and a colonoscopy was repeated at week 16. In some patients, due to the Covid-19 pandemic, technical or personal issues, follow-up endoscopies were conducted at week 24 or 48 instead.

### Assessment of disease activity

Response was defined as a decrease in the endoscopic Mayo score of at least 1 point from baseline and/or an objective improvement in disease severity (e.g., a decrease in endoscopic disease extension -number of segments with an endoscopic Mayo score >1) or a partial Mayo score ≤ 2 with no item >1). Histologic evaluations were performed by an expert pathologist (MR) in a blinded fashion as described below. Table 1 shows the demographic and clinical characteristics at baseline and during follow-up for responders (R) and non-responders (NR).

### Sample collection

Endoscopic colonic biopsies (4-8 from each patient) were collected from the involved sigmoid colon (or rectum in the case of isolated proctitis) at baseline (week 0) and from the same area (irrespective of disease activity) during follow-up. Serum and whole-blood RNA (Paxgene tubes, BD Biosicences) were also collected at baseline, as well as at weeks 2, 8 and 16, when possible; otherwise at week 24 or 48. In addition, biopsies from healthy non-IBD individuals (n=10) were obtained to serve as healthy controls (HCs) for bulk RNA analysis (male:female=6:4; median age and range=46 (38–70)).

### Human colonic cell isolation for single-cell RNA-seq analysis

To generate single-cell suspensions, colonic biopsies (n=3-4 per patient) were enzymatically digested using 0.5 Wünsch units/ml Liberase TM (Roche) and DNase I at 10µg/ml at 37°C (a protocol previously optimized in our lab)(43). Digested samples were filtered through both 50µm and 30µm filters before processing for scRNA-seq analysis.

### Single-cell RNA sequencing of intestinal cells

Cell suspensions were processed using the 10× Genomics 3′ mRNA single-cell method and loaded onto the Chromium10x Genomics platform (10x Genomics, CA, USA). Approximately 7,000 cells were loaded onto the Chromium10x Genomics platform (10x Genomics, CA, USA) following the manufacturer’s protocol. The generation of gel beads in emulsion (GEMs) (10x Genomics, CA, EEU), barcoding and GEM-reverse transcription were performed using the Chromium Single Cell 3′ and Chromium Single Cell V(D)J Reagent Kits (10× Genomics, CA, EEU) (user guide, no. CG000086) according to the manufacturer’s instructions. Full-length, barcoded cDNA was amplified by PCR to generate enough mass for library construction (Nextera® PCR primers) (Illumina, CA, USA). Sequencing of the libraries was performed on HiSeq2500 (Illumina, CA, USA).

### Single-cell data analysis

For each sample, sequences obtained in fastq files were processed with a CellRanger’s count pipeline (10XGenomics, v.3.1.0) using the default parameters. This pipeline performs an alignment based on the reference genome (Gencode release 27, assembly GRCh38 p10), followed by filtering, barcode counting, and UMI counting. The resulting filtered matrix was analyzed using R (v.4.1.2). We merged the count matrices retrieved from CellRanger using the function *merge* from the SeuratObject R package (version 4.0.2). At this point, doublets were assessed using the scDblFinder R package (version 3.1.0) and removed. Once we had all of the samples in the same object, we filtered the low-quality cells based on the mitochondrial RNA percentage and the number of genes per cell. A total of 69,813 high-quality single cells were considered for the analysis. We then logarithmically normalized them, obtained the highly variable genes, and scaled the counts (default parameters) of each data set using Seurat (version 4.1.0). Principal component analysis (PCA) was subsequently performed. Dimensionality reduction was accomplished by applying the Uniform Manifold Approximation and Projection (UMAP) algorithm using the optimal number of principal components. UMAP also served as a two-dimensional embedding platform for data visualization. Cluster analysis was performed using the Louvain clustering algorithm. Based on the marker genes of the clusters (Wilcoxon test against all the other clusters, false discovery rate adjusted p.value < 0.05), cells were initially classified into 6 groups (epithelium, T cells, plasma and B cells, myeloid cells, stroma and cycling cells). Whereas most subsets were filtered using 25% mitochondrial gene expression, epithelial cells required a less stringent filter of 65%. We then systematically re-clustered each cell category using an unsupervised Louvain clustering algorithm. The annotation of each subcluster from the main cell type was defined by the marker genes, which were obtained by the *FindAllMarkers* function with the default threshold, except for the *min.pct* parameter, which was set to 0.25, and the *thresh.use*, which was set to 0.25.

### Identification of cell types

Each main cell type was re-processed starting from *FindVariableFeatures* using the same procedure as for the whole dataset (see the full code at this <u>link</u>). During this process, doublets were identified through expert annotation of the marker gene lists for each cell cluster and matched to clusters with markers from distinct lineages. We then systematically re-clustered each cell category using an unsupervised Louvain clustering algorithm. The annotation of each subcluster from the main cell type was defined by the marker genes, which were obtained by the *FindAllMarkers* function with the default threshold, except for the *min.pct* parameter, which was set to 0.25, and the *thresh.use*, which was set to 0.25.

### Batch correction

To correct batch effects across samples, we applied Harmony (version 0.1.0). Specifically, we used the *RunHarmony* function, with the optimal number of principal components found for each subset as latent space and the sample of origin as the batch label. Seurat’s analysis pipeline was applied again to each object, followed by UMAP generation using default settings and harmony integrated space.

### Annotation of cells

Markers defining each cell type are summarize in Supplementary Table 4 and Supplementary table 5.

### Cell communication analysis

For cellular communication analysis, we used CellphoneDB (version 4.1.0) across the distinct cell types defined. For each comparison, we used the normalized counts and calculated the average expression levels of the interacting molecules in both the ligand and receptor populations.

### Pathway activity inference

We used PROGENy (8) (version 1.26.0) to infer pathway activities of NFkB and JAK-STAT into our previously curated Seurat object. We computed the mean activity of each cell type and then performed comparisons at baseline (R Pre-tx vs NR Pre-tx) and during treatment (R Pre-tx vs R Post-tx, and NR Pre-tx vs NR Post-tx) using a Wilcoxon signed-rank test adjusted with a Bonferroni correction.

### Population similarity analysis

Statistically significant marker genes from the INHBA+ and FOLR2+ macrophage populations were identified using *FindAllMarkers.* Then, matchSCore R package(44) was used to perform Jaccard similarity indexing between the different clusters and the markers for M1 and M2 from Garrido-Trigo *et al* (43).

### RNA-seq analysis

Raw RNA-seq data from GSE181250 (11) was analyzed, and genes with low expression levels and low variations across the samples were removed. Differentially expressed genes between the LPS-stimulated macrophages treated with a blocking anti-IL-10 antibody and control groups were identified using DESeq2(45).

### Pathway analysis

Differentially down-regulated genes (FC < 0.86 & fdr < 0.05) in the non-responder S1 population were identified using the *FindMarkers* function in the Seurat package(46). Gene Ontology (GO) enrichment analyses for biological processes were conducted using the *enrichGO* function in the clusterProfiler package(47).

### Macrophage differentiation

Human peripheral blood mononuclear cells (PBMCs) were isolated from buffy coats of healthy volunteers over a Lymphoprep (Nycomed Pharma) gradient according to standard procedures. Monocytes were purified from PBMCs by magnetic cell sorting using CD14 Microbeads (Miltenyi Biotec). Monocytes (> 95% CD14+ cells) were cultured at 0.5 × 106 cells/ml for 7 days in RPMI medium supplemented with 10% fetal calf serum at 37°C in a humidified atmosphere with 5% CO2. Monocyte colony-stimulating factor (M-CSF; 10 ng/ml; ImmunoTools) was added at days 0, 2 and 5. M-CSF was chosen as the differentiation model due to its ability to generate macrophages that exhibit the highest similarity to the FOLR2+ macrophages identified in our dataset(^43, 48^).

### Isolation and culture of intestinal fibroblasts

Fibroblasts were isolated from the non-tumoral tissue of colorectal cancer patients undergoing surgery. Tissue (2 cm^2^) was first incubated with 1 mM DTT and 5 mM EDTA in 1xHank’s balanced salt solution (HBSS), followed by an incubation with 5 mM EDTA in HBSS, both for 30 min at 37 °C with agitation. Tissue was then minced and digested with 5.4 U/mL collagenase D (Roche Applied Science), 100 U/mL DNase I (Sigma), 39.6 U/mL dispase II (Gibco) and 1% of antibiotic solution in Dulbecco’s Modified Eagle Medium (DMEM) for 20 minutes at 37 °C with agitation. The cell suspension was filtered and centrifuged at 500 *g* for 5 min. After being treated with Red Blood Cell Lysis Buffer (Biolegend), the single-cell suspension was cultured in a T-25 flask (TPP) with RPMI1640 medium (Gibco), supplemented with 10% FBS (VWR), 1% HEPES (Gibco), 1% L-glutamine (Gibco), 1% sodium pyruvate (Gibco), and 1% antibiotic/antimycotic solution (Sigma-Aldrich). The next day non-attached cells were removed. When fibroblasts reached confluency, cells were passaged using 0.25% Trypsin-EDTA (Gibco).

### Macrophage and fibroblast activation and treatment with tofacitinib

On day 6 of macrophage differentiation or during passage 2-5 of the fibroblasts culture, tofacitinib (or vehicle control) was added at a final concentration of 300 nM, followed by inflammatory stimuli tumor necrosis factor (TNF; 20 ng/ml), lipopolysaccharide (LPS; 10 ng/ml) or interferon-gamma (IFNγ; 5 ng/ml) 30 min later. Cells were incubated at 37 °C for 19 h. Cells and supernatants were collected for Real-time PCR (qPCR) and enzyme-linked immunosorbent assay (ELISA) analysis, respectively.

### RNA isolation

Biopsies (n= 10 from non-IBD controls, n= 29 biopsies from 15 UC patients) were placed in RNA later (Qiagen) and stored at −80°C until further analysis. Blood (n= 10 non-IBD controls, n= 48 samples from 15 UC patients) was collected in PAXgene Blood RNA collection tubes (PreAnalytiX) and stored at −20°C. Total RNA was extracted using a RNeasy Kit (Qiagen) according to the manufacturer’s instructions.

Macrophages and fibroblast were harvested, and RNA was isolated using a Mini Kit for RNA and protein purification (NucleoSpin RNA/Protein, Macherey-Nagel) according to the manufacturer’s instructions.

For all sample types, the purity and integrity of the total RNA were assessed using the 2100 Bioanalyzer (Agilent) and then quantified via a NanoDrop spectrophotometer (Nanodrop technologies); only samples with an RNA integrity number (RIN) >7.0 were used.

### Quantification of gene expression (qPCR)

Biopsy, blood, macrophage and fibroblast RNA was retro-transcribed to cDNA using a reverse transcriptase (High-Capacity cDNA Archive RT kit; Applied Biosystems). qPCR analysis was performed in an ABI PRISM 7500 Fast RT-PCR System (Applied Biosystems) or QuantStudio 3 using TaqMan Universal PCR Master Mix (Applied Biosystems) and predesigned primers and a probe specific for each target of interest (Supplementary Table 6). ACTB was used as the endogenous control (TaqMan primers and probes; Applied Biosystems). Data are expressed as arbitrary units (AU), applying the 2^-dCt^ formula where the dCt is the difference between the mean Ct of the target gene and the mean Ct of the endogenous gene.

### Cytokine measurement by an enzyme-linked immunosorbent assay (ELISA)

Cytokine detection was carried out in cultured macrophage and fibroblast supernatants using ELISA kits for human IL-10, IL-6, TNF, CXCL10 and CXCL1 (DuoSet^TM^ ELISA, R&D Systems) according to the manufacturer’s instructions.

### H&E and semi-quantitative histological scores

Conventional staining with Hematoxylin Eosin (H&E) was performed with the Stainer Integrated Workstation Leica ST5010-CV5030 (Leica Biosystems, Nussloch, Germany) on 3-micron tissue sections placed on microscope slides according to the manufacturer’s protocol. All slides were scanned at low power with an optical microscope (Olympus BX41, Olympus, Japan) and inflammatory cell visualization was performed using the 20-objective lens corresponding to an area of 0.785 mm^2^.

A pathologist blindly scored the severity of histologic inflammation, categorizing it as non-inflamed (score=0), mildly (score=1), moderately (score=2), or severely (score=3) inflamed. A pathologist also semi-quantified the amount of plasma cells, lymphocytes, neutrophils, eosinophils, endothelium, fibroblasts, and epithelium present in each H&E –stained slide over a range from 0 to 3, which we arbitrarily defined with a score of 0. This reflected the amount of cells present in a healthy intestinal mucosa and the maximum value of 3, which indicated the highest abundance of cells in the study cohort. In the context of the epithelium, a score of 0 indicates a complete epithelial loss and a score of 3 signifies a healthy epithelial layer.

### Measurement of serum tofacitinib concentration (Barcelona cohort only)

Serum tofacitinib concentrations were measured at the Unitat de Tècniques Separatives (CCiT, University of Barcelona, Spain). Tofacitinib (Tofacitinib citrate PZ0017, Sigma) standard was prepared using 100 μL of serum (without tofacitinib), 20 μL of 100ng/ml internal standard (IS; [13C3, 15N]-Tofacitinib C3928, Alsachim), and 20 μL of standard working solution prepared via a serial dilution method of Tofacitinib in methanol (0-0.1-0.5-2-10-25-50 ng/ml) and 300 μL of acetonitrile.

For patient sample preparations, 100 μL of serum were spiked with 20 μL of IS 100 ng/ml, 20 µl methanol and 300 μL of acetonitrile. Both standards and samples were vortexed for 1 min and centrifuged at 13000 × *g* for 10 min at 4 °C. Supernatants were analyzed using a High-performance Liquid Chromatography (HPLC) MS-MS system. The HPLC Acquity (Waters) was used in combination with a 4000QTRAP system from Sciex (Sciex, Foster City, CA, USA) with a TurboV Ion Source operating in positive ion mode. The chromatographic separation was achieved on a Kinetex 2.6µm PFP 50×2.1 mm column (Phenomenex, Torrance, CA, USA). The mobile phase was composed of 10mM NH_4_HCOO pH 4.1 (eluent A) and acetonitrile (eluent B) at a flow rate of 0.5 ml/min and using the following gradients (t(min), %A): (0, 5), (12, 95), (14, 95), (14.5, 5), (17, 5). The injection volume was set at 2 μL. Detection using mass spectrometry was carried out in multiple reaction monitoring (MRM) mode with the transitions set as follows: 313.1/149.1 (identification and quantitation), 313.1/98.2 (confirmation) for tofacitinib and 317.7/149.1 for the IS.

### Statistical analysis

Graphs were generated using R (version 4.3.2) and Graphpad Prism (Graphpad Software, version 10). The difference between tofacitinib serum concentrations in the R and NR groups was tested using the Wilcoxon signed-rank test. Statistical differences in single-cell RNA sequencing (scRNA-seq) population cell abundances were assessed using the chi-squared contingency table test (*chisq.test* function in R stats), considering only those differences with a fold change > 1.5 or < 0.66. qPCR fold-change results were log2-transformed and subjected to a one-sample t-test. ELISA results were analyzed using a paired t-test. Both analyses were executed through the t.test function in R stats. Differences in patient characteristic were assessed using the Fisher’s exact test (*fisher.test* function in R) for categorical variables and the Wilcoxon rank-sum test for numerical variables. Differences in blood populations and gene expression quantification were evaluated using the Kruskal-Wallis rank-sum test (*kruskal.test*). To account for multiple comparisons, a false discovery rate (FDR) control was implemented, and corrected p-values < 0.05 were deemed statistically significant.

### Study approval

The study was approved by the Hospital Clinic Barcelona Ethics Committee (HCB/2019/0714), by the Departament de Salut of the Generalitat de Catalunya (0336/16898/2019), by the Agencia Española de Medicamentos y Productos Sanitarios (AEMPS) and by the UZ Leuven Institutional Review Board (BD Biobank CCARE B322201213950/S53684/S68081). All patients signed an informed consent form prior to enrolling in the study.

## Data availability

The raw data from single-cell RNA sequencing and all source code will be available in the Gene Expression Omnibus (GEO) and on GitHub, respectively, upon acceptance of this paper.

## Abbreviations

BID: "bis in die" twice a day
DEG: differentially expressed genes
ELISA: enzyme-linked immunosorbent assay
GM-CSF: granulocyte-monocyte colony-stimulating factor
HC: healthy control
IBD: inflammatory bowel disease
IFN: interferon
IL: interleukin
JAK: Janus kinases
LPS: lipopolysaccharide
M-CSF: macrophage colony-stimulating factor
NR: non-responders
qPCR: Real-time PCR
RNA: ribonucleic acid
scRNA-seq: single-cell RNA sequencing
TNF: Tumor necrosis factor
UC: ulcerative colitis

## Writing assistance

Writing assistance was funded by a Research grant from Pfizer.

## Author contributions

EM-A: Data acquisition and curation, investigation, methodology, writing – review & editing. MV: Data curation, methodology, writing – review & editing. AMC: Data curation, formal analysis, software, visualization. AG-T: Investigation, methodology. VG: Investigation, methodology, writing – review & editing. AS-M: Data curation, formal analysis, software, visualization. MB: Data curation, formal analysis, software, visualization. ME: Investigation. MR: Methodology, investigation. MCM: Investigation, resources. AG: Investigation, resources. IO: Investigation, resources. AF-C: Investigation, resources. BC: Investigation, resources. AC: Methodology, writing – review & editing. BV: Investigation, resources, data curation, writing – review & editing. SV: Investigation, Resources, data curation, writing – review & editing. JP: Conceptualization, funding acquisition, writing – review & editing. ER: Investigation, resources, writing – review & editing. AS: Conceptualization, project administration, writing – original draft, data curation, funding acquisition.

## Acknowledgements

We thank Dr. Amaya Puig-Kruger at Hospital General Universitario Gregorio Marañon, Madrid, Spain for her support in setting up experiments with monocyte-derived macrophages. We thank the Single Cell Genomics group at CNAG (Barcelona, Spain) and the Unitat de Tècniques Separatives at CCiT, University of Barcelona (Spain) for their technical support. This work was supported by Pfizer and Grant PID2021-123918OB-100 from the Ministerio de Ciencia e Innovación, Gobierno de España. EM-A was funded by fellowship RH042155 (RTI2018 096946-B-I00) from the Ministerio de Ciencia e Innovación, Gobierno de España.

## Supplementary Material

**Melón-Ardanaz E *et al*.**

**Supplementary Table 1.** Summary of demographic and clinical characteristics of study participants from both recruiting sites.

| **Characteristics at baseline** | **Barcelona** | **Leuven** |
| --- | --- | --- |
| N | 17 | 14 |
| Sex (male/female) | 8/7 | 9/5 |
| Age (median, range, years) | 41 (22-55) | 47.1 (18.1-73.6) |
| Disease duration (median, range, years) | 8 (2-28) | 11.1 (2-27.5) |
| UC extension (Pancolitis/Left-sided colitis/Proctitis) | 5/7/05 | 6/7/01 |
| **Current treatment** |  |  |
| Aminosalicylates (e.g., mesalazina) | 3 | 5 |
| Corticosteroids (e.g., beclometasona) | 6 | 0* |
| Immunomodulator (e.g., Azathioprine) | 6 | 0* |
| Monotherapy (one of the above) | 10 | 5 |
| Combination therapy (two or more of the above) | 3 | 0 |
| Number of previous biologics (median, range) | 2 (0-4) | 2.5 (2-3)* |
| CRP (mean, SD) | 0.63 (0.43) | 0.84 (0.98) |
| Endoscopic Mayo (median, range) | 3 (2-3) | 3 (2-3) |
| Total Mayo (median, range) | 8 (4-11) | 10.5 (8-12)* |

\* p-value <0.05

**Supplementary Table 4.**
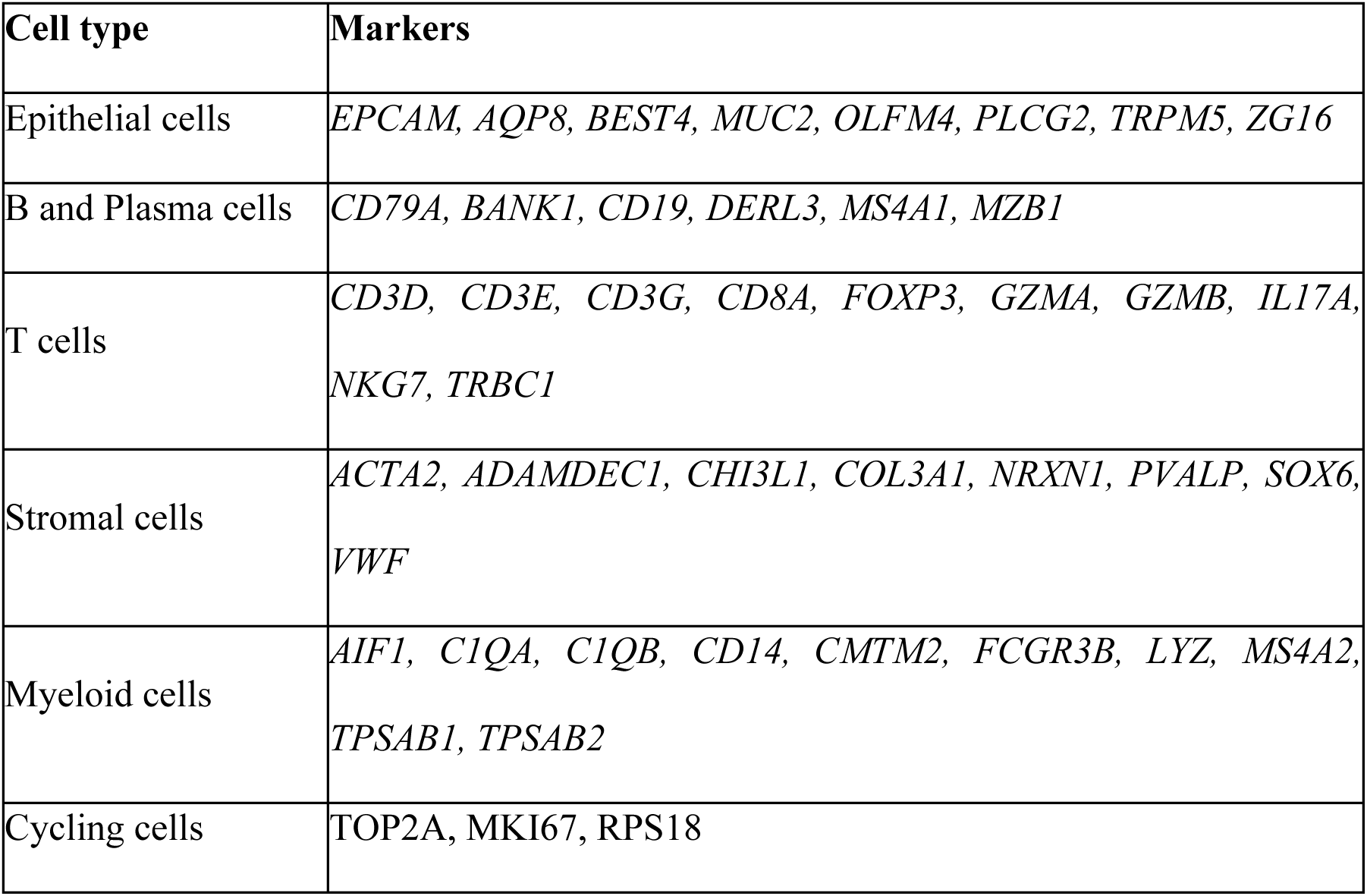
Gene markers used to separate each of the six main cell types identified by scRNA-seq.

**Supplementary Table 5.**
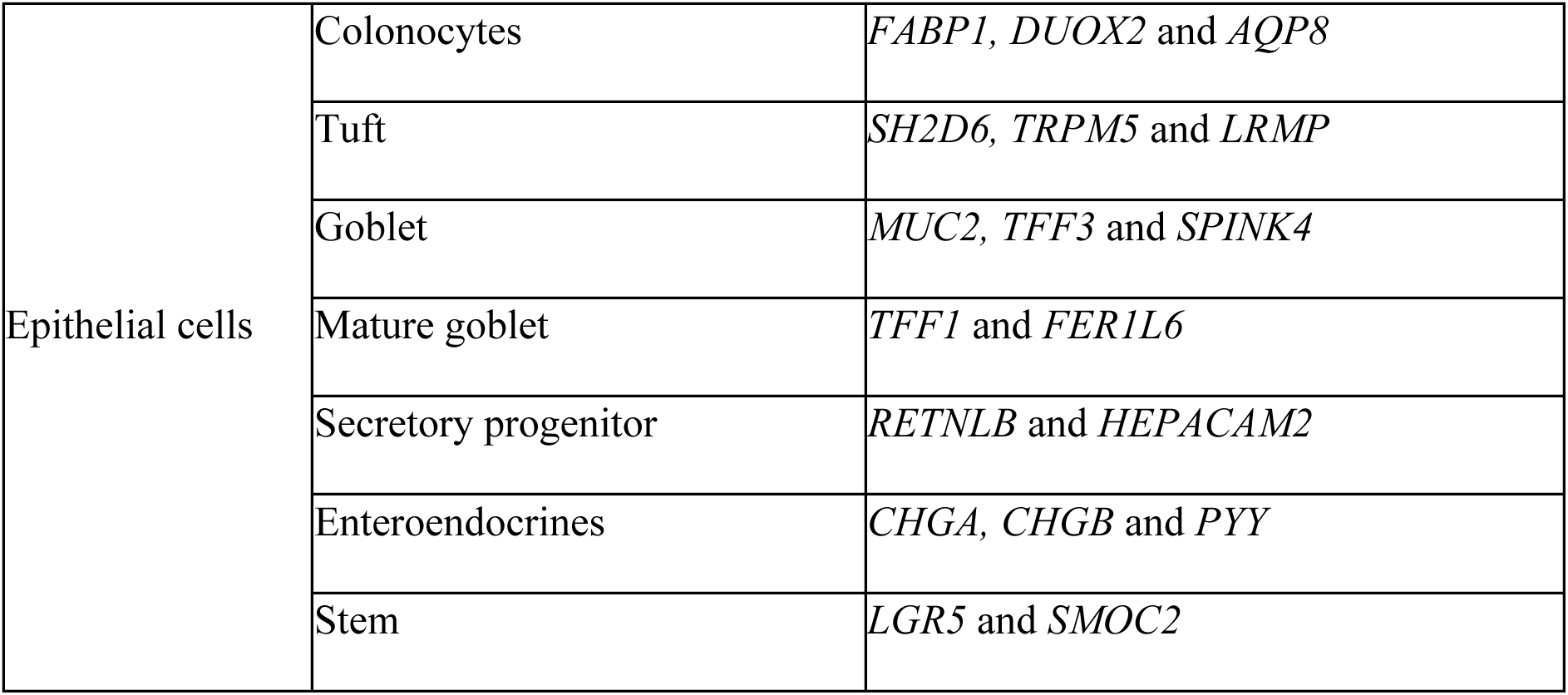
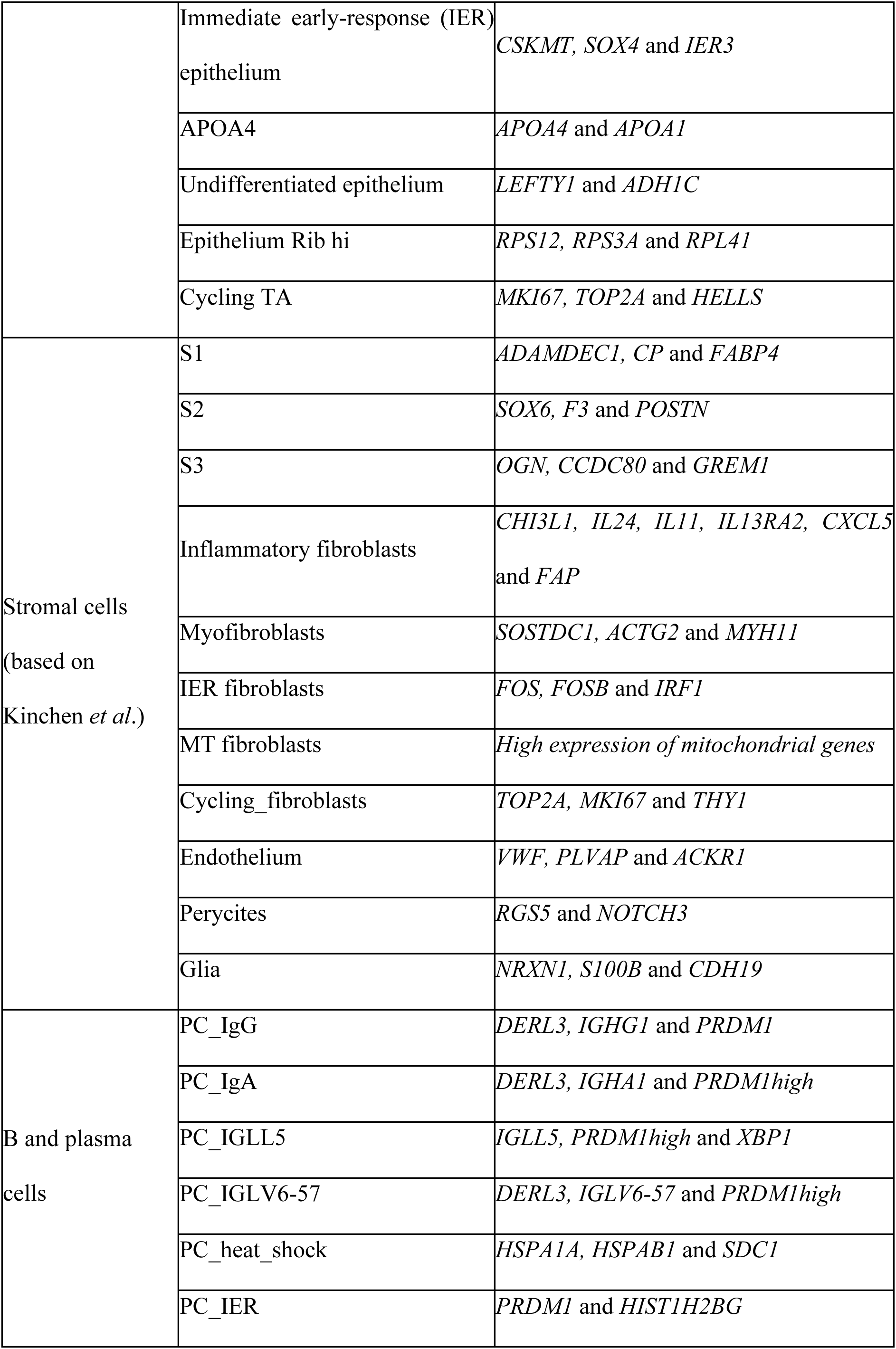
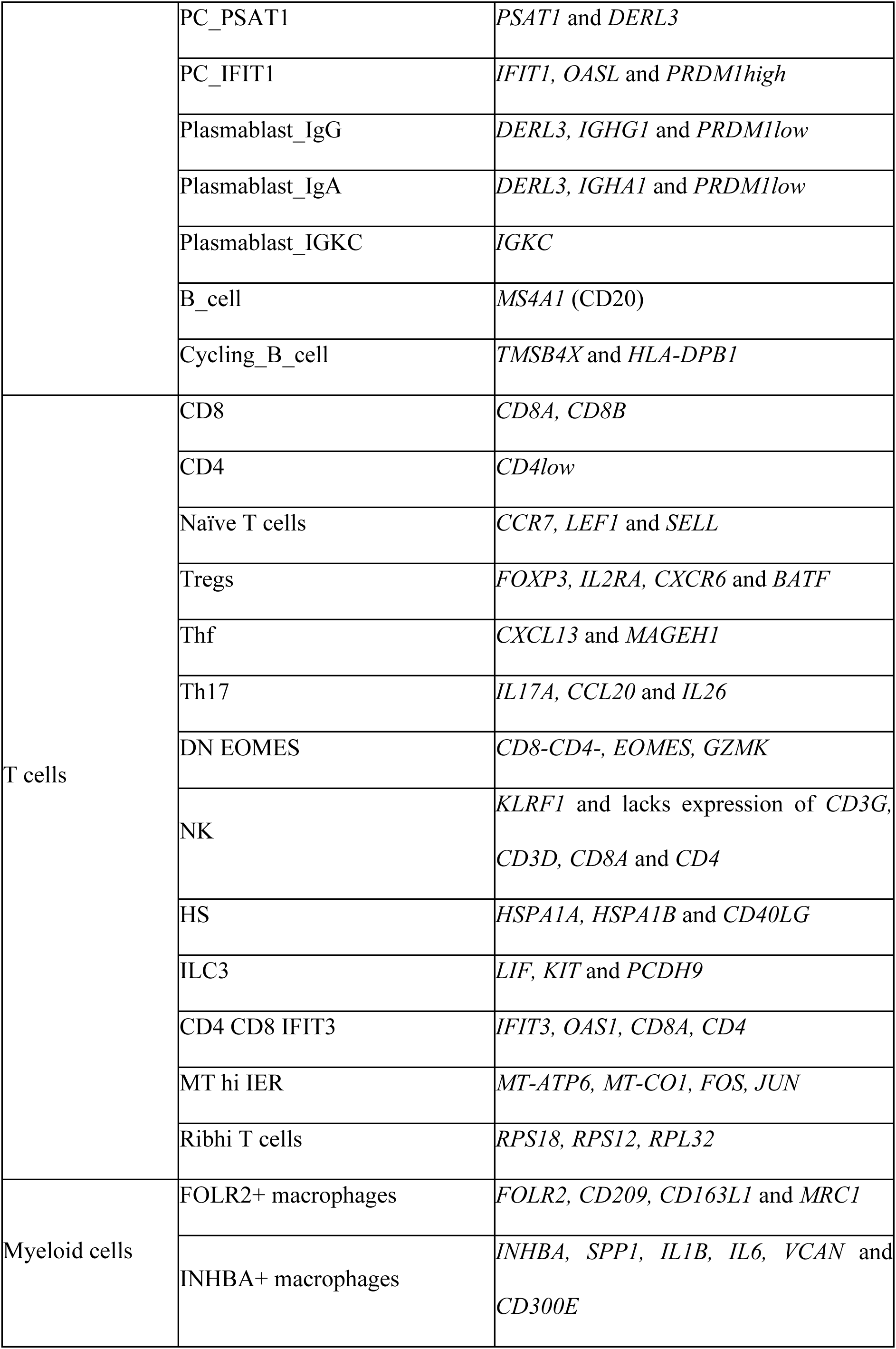
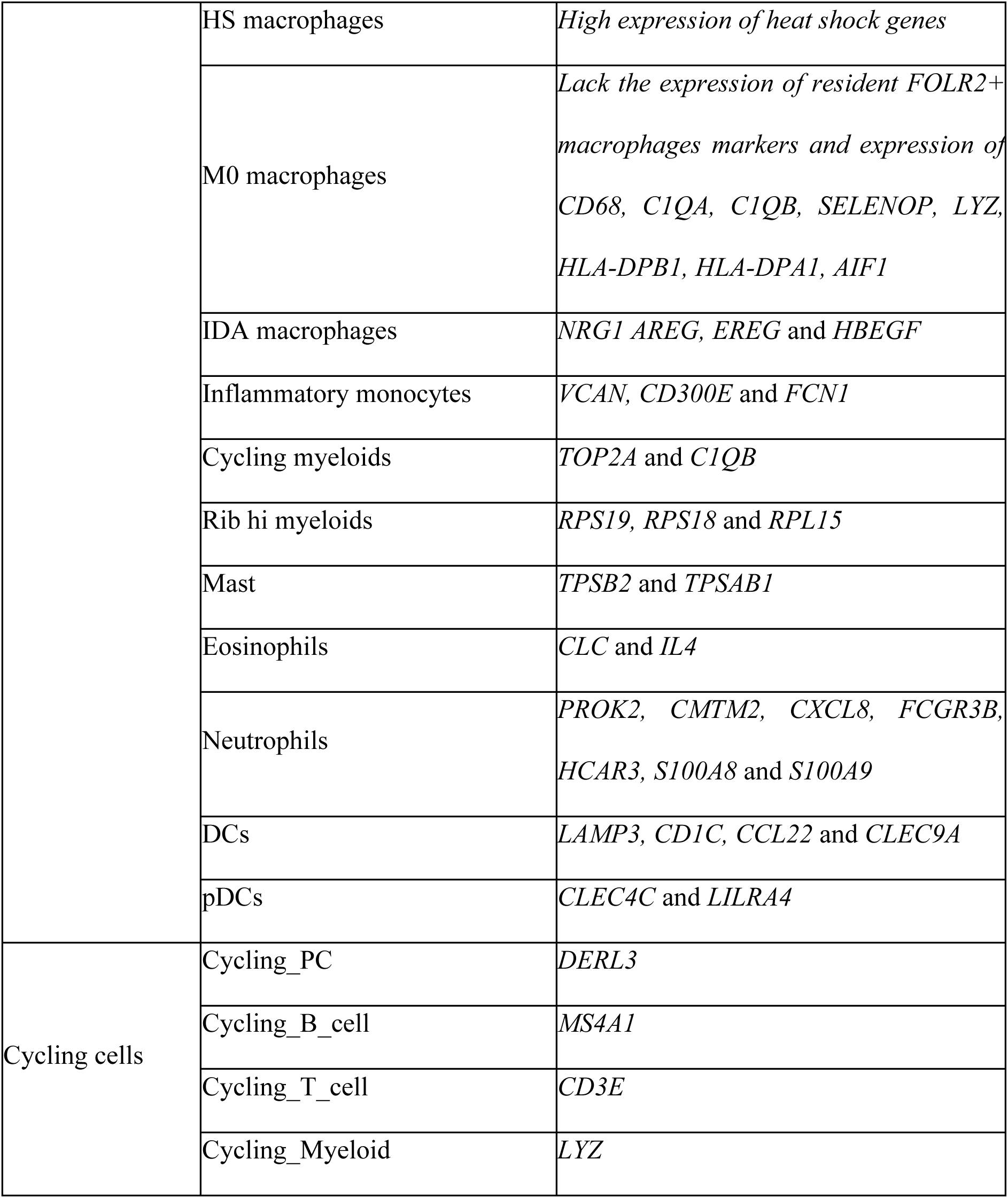
Gene markers used to annotate all of the clusters within each of the six main cell types identified by scRNA-seq.

**Supplementary Table 6.** TaqMan primers probe references.

| Gene | Reference |
| --- | --- |
| ACTB | Hs99999903_m1 |
| ACOD1 | Hs00985781_m1 |
| AQP8 | Hs00154124_m1 |
| CCL5 | Hs00982282_m1 |
| CD209 | Hs01588349_m1 |
| CD3E | Hs01062241_m1 |
| CDH1 | Hs01023894_m1 |
| CHI3L1 | Hs01072228_m1 |
| CTLA4 | Hs00175480_m1 |
| CXCL10 | Hs01124251_g1 |
| CXCL5 | Hs01099660_g1 |
| CXCL8 | Hs00174103_m1 |
| CLEC5A | Hs04398399_m1 |
| DERL3 | Hs00405322_m1 |
| MS4A1 | Hs00544819_m1 |
| EPCAM | Hs00901885_m1 |
| FAP | Hs00990791_m1 |
| FCGR3A | Hs02388314_m1 |
| FOXP3 | Hs01085832_m1 |
| IDO1 | Hs00984148_m1 |
| IL1B | Hs01555410_m1 |
| IL10 | Hs00174086_m1 |
| IL23A | Hs00372324_m1 |
| IL6 | Hs00985639_m1 |
| INHBA | Hs01081598_m1 |
| IRF1 | Hs00971965_m1 |
| LCN2 | Hs01008571_m1 |
| LTB | Hs00242739_m1 |
| MMP9 | Hs00234579_m1 |
| MX1 | Hs00895608_m1 |
| OAS1 | Hs00973635_m1 |
| PDGFRA | Hs00998018_m1 |
| PROK2 | Hs00363716_m1 |
| S100A8 | Hs00374264_g1 |
| SOCS1 | Hs00705164_s1 |
| SOCS3 | Hs02330328_s1 |
| SPP1 | Hs00959010_m1 |
| TFF3 | Hs00902278_m1 |
| TNF | Hs01113624_g1 |
| WNT5A | Hs00998537_m1 |

## Supplementary Figures

**Supplementary Figure 1.**
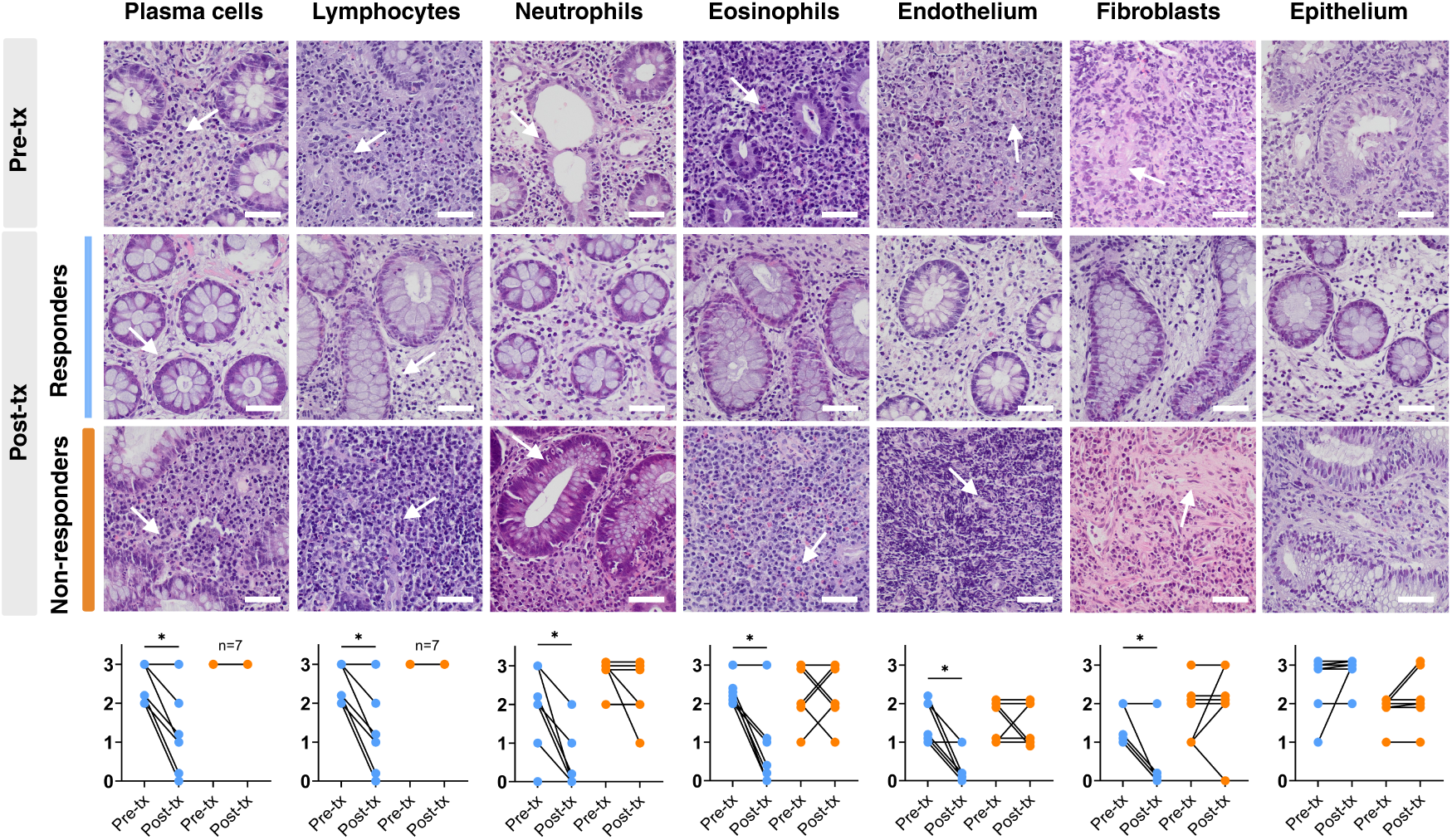
Evaluation of histologic disease activity in patients treated with tofacitinib. Representative hematoxylin and eosin staining of colonic samples from UC patients pre- and post-tofacitinib treatment. Seven colonic cell types (examples shown with white arrows) could be distinguished by an expert pathologist (MR). Scale bar 20µm. Semi-quantitative scores (see Supplementary Methods) for the abundance of each cell type in UC patients pre- and post-tofacitinib treatment in responders (n= 7) and non-responders (n= 7). Wilcoxon test for matched data (two-tailed p-value). *p<0.05.

**Supplementary Figure 2.**
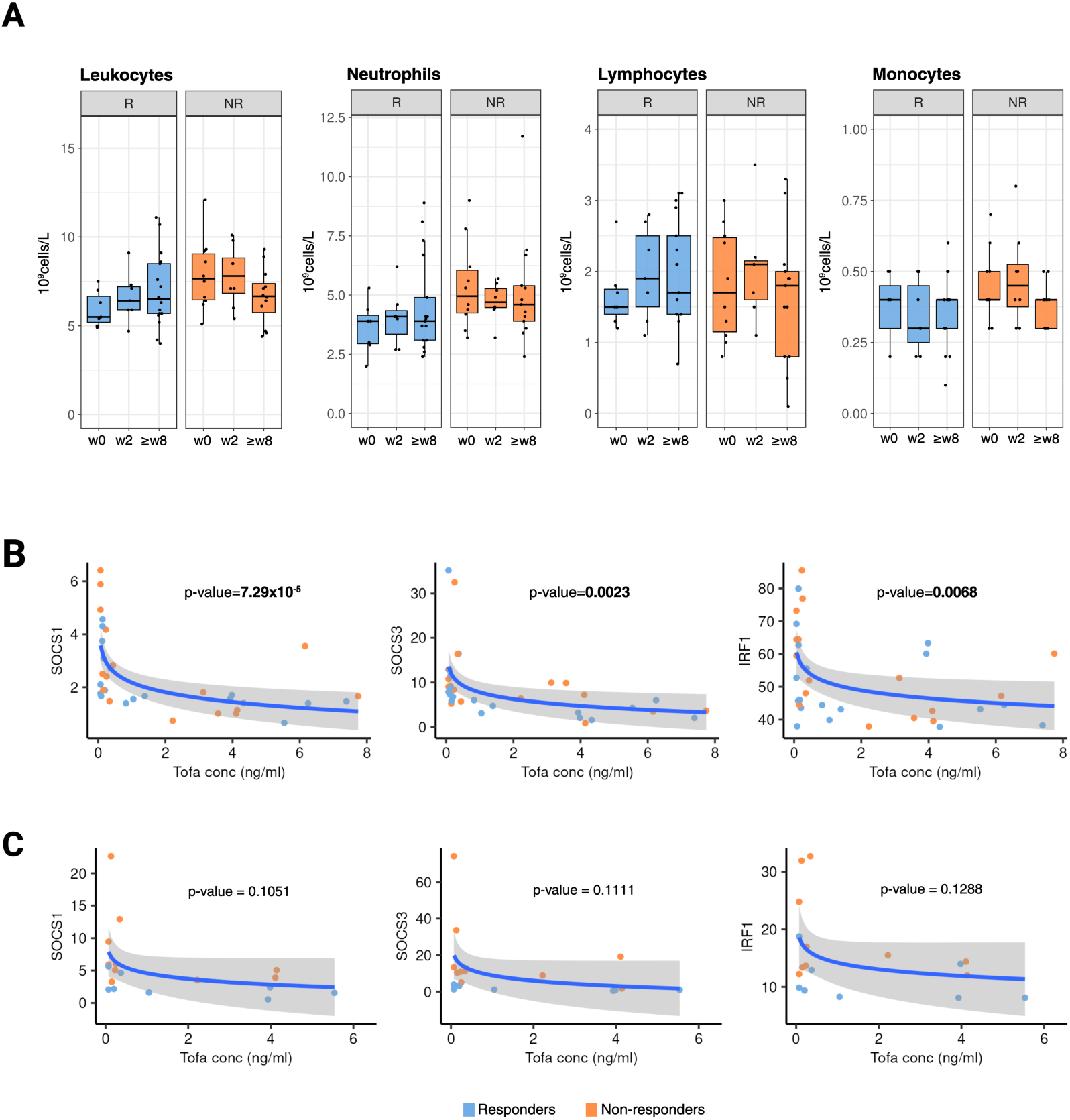
Effects of tofacitinib on blood leukocytes and on blood and mucosal gene expression. A) Concentrations (10^9^cells/L) of total leukocytes, neutrophils, lymphocytes, and monocytes in the peripheral blood of tofacitinib-treated patients split by responders (R; blue) and non-responders (NR; orange). Data is shown at baseline (w0; R n=7 and NR n=10) and at early (w2; R n=7 and NR n=9) and later (≥8w; R n=17 and NR n=13) follow-up timepoints. Data are presented as boxplots. No comparison was found significant; Kruskal-Wallis rank-sum test. B) Correlation analysis of tofacitinib serum concentrations and mRNA expression (expressed as arbitrary units) of relevant JAK-STAT pathway target genes in the blood (n=37) or C) biopsies (n=18) of tofacitinib-treated patients. Log-linear regression analysis. Responders and non-responders are shown in blue and orange, respectively.

**Supplementary Figure 3.**
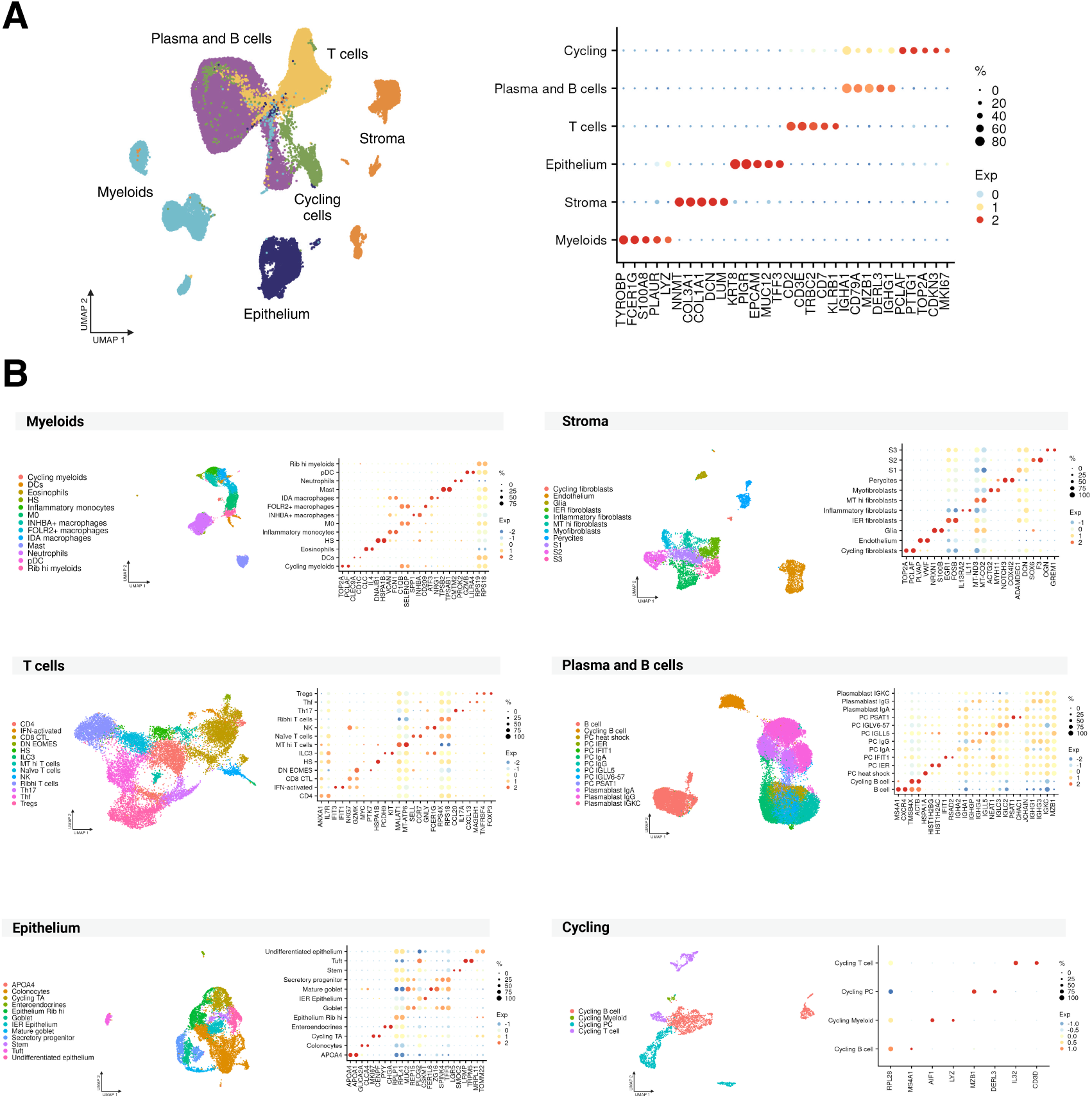
Annotation of cell types in intestinal single-cell RNA sequencing data. Uniform manifold approximation and projection (UMAP) of scRNA-seq data for (A) each main cell type and (B) further sub-clustering of each cellular type, colored by the different cell subsets and dot plots showing the fraction of expressing cells (dot size) and mean expression levels (dot color) of the principal markers for each of these subpopulations.

**Supplementary Figure 4.**
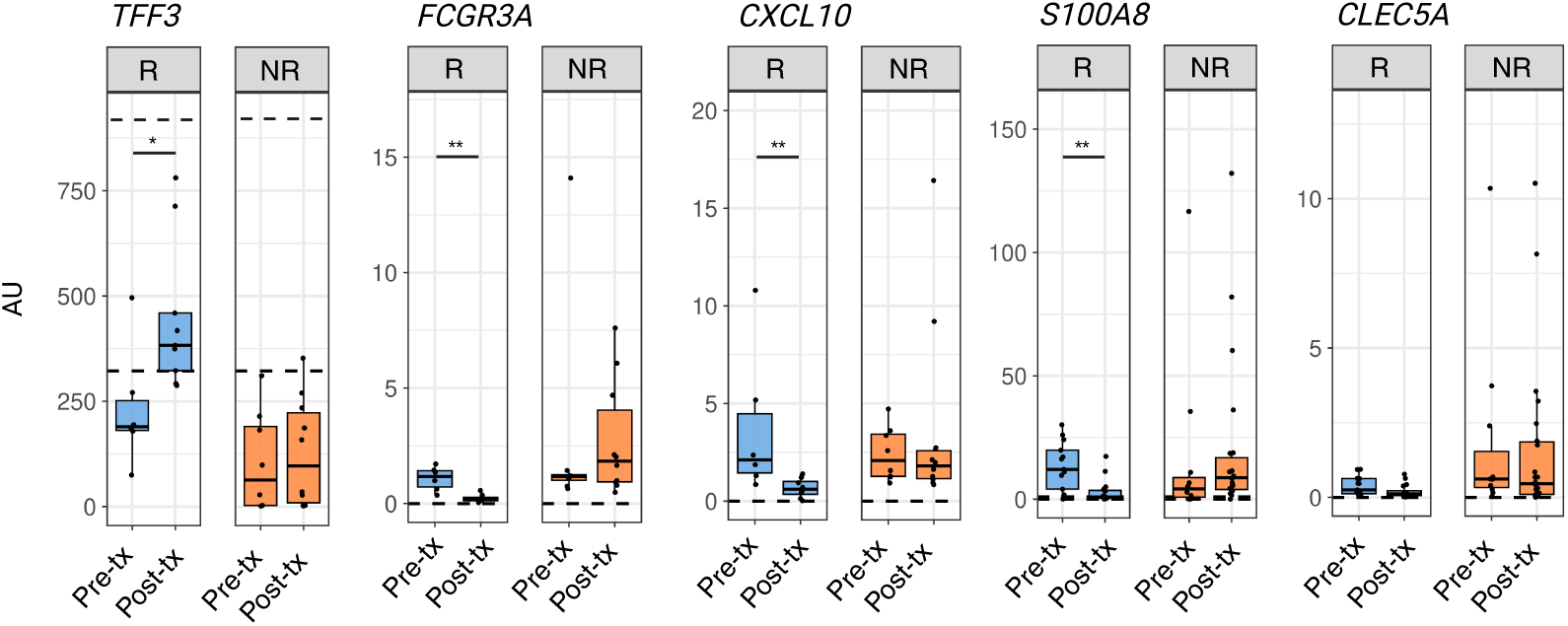
Expression of selected genes in bulk RNA from intestinal biopsies of patients receiving tofacitinib. Gene expression of whole biopsy bulk RNA for additional markers of cellular subsets including *TFF3* (goblet cells), *FCGR3A* (monocytes and macrophages), inflammatory markers (*CXCL10* and S100A8) and *CLEC5A* (INHBA+ macrophages). Data is shown separately for responders (R; blue) pre-treatment (pre-tx, n=13 samples) and post-treatment (post-tx, n=17 samples); and non-responders (NR; orange) pre-tx (n=11 samples) and post-treatment (n = 22 samples). Data is expressed as arbitrary units (AU). Dotted lines show the standard error of the mean of biopsies from healthy controls (n=10). Groups were compared using a Wilcoxon test and corrected for multiple testing. Data are expressed as median±range. False discovery rate-corrected p-values: *< 0.05 and **< 0.01.

**Supplementary Figure 5.**
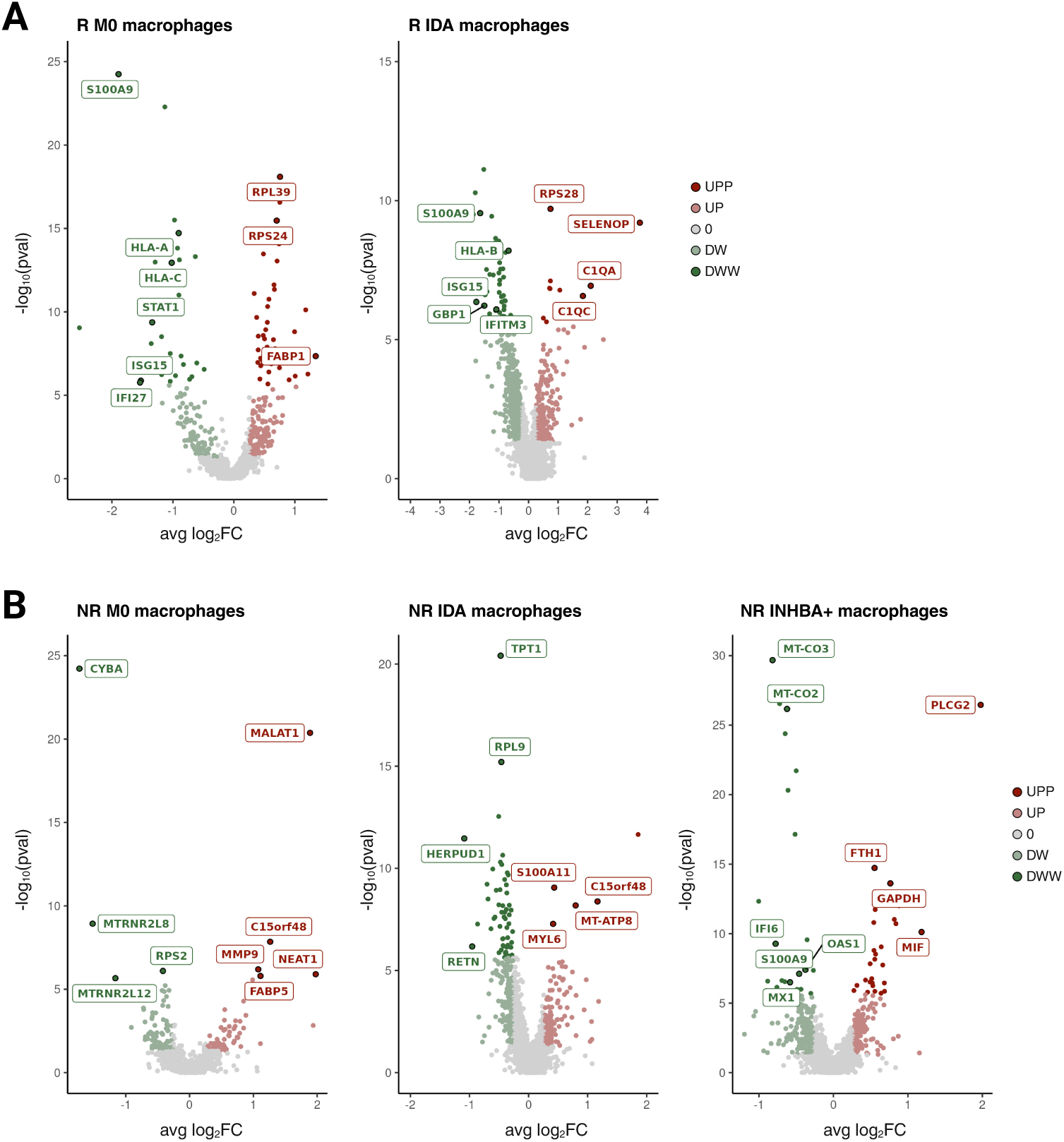
Tofacitinib induces transcriptional changes in subsets of intestinal macrophages. Volcano plots showing the principal differentially expressed genes during treatment (post-tx vs pre-tx) in the macrophage subsets of responders (A) and non-responders (B). A two-sided Wilcoxon rank sum test was applied. Genes with a false discovery rate adjusted p value < 0.05, and a fold change (FC) > 1.2 (UPP, dark red) or FC < 0.83 (DWW, dark green) were considered regulated. Additionally, genes with a nominal p-value < 0.05 are shown in light red (UP) if FC > 1.2 and light green (DW) if FC < 0.83.

**Supplementary Figure 6.**
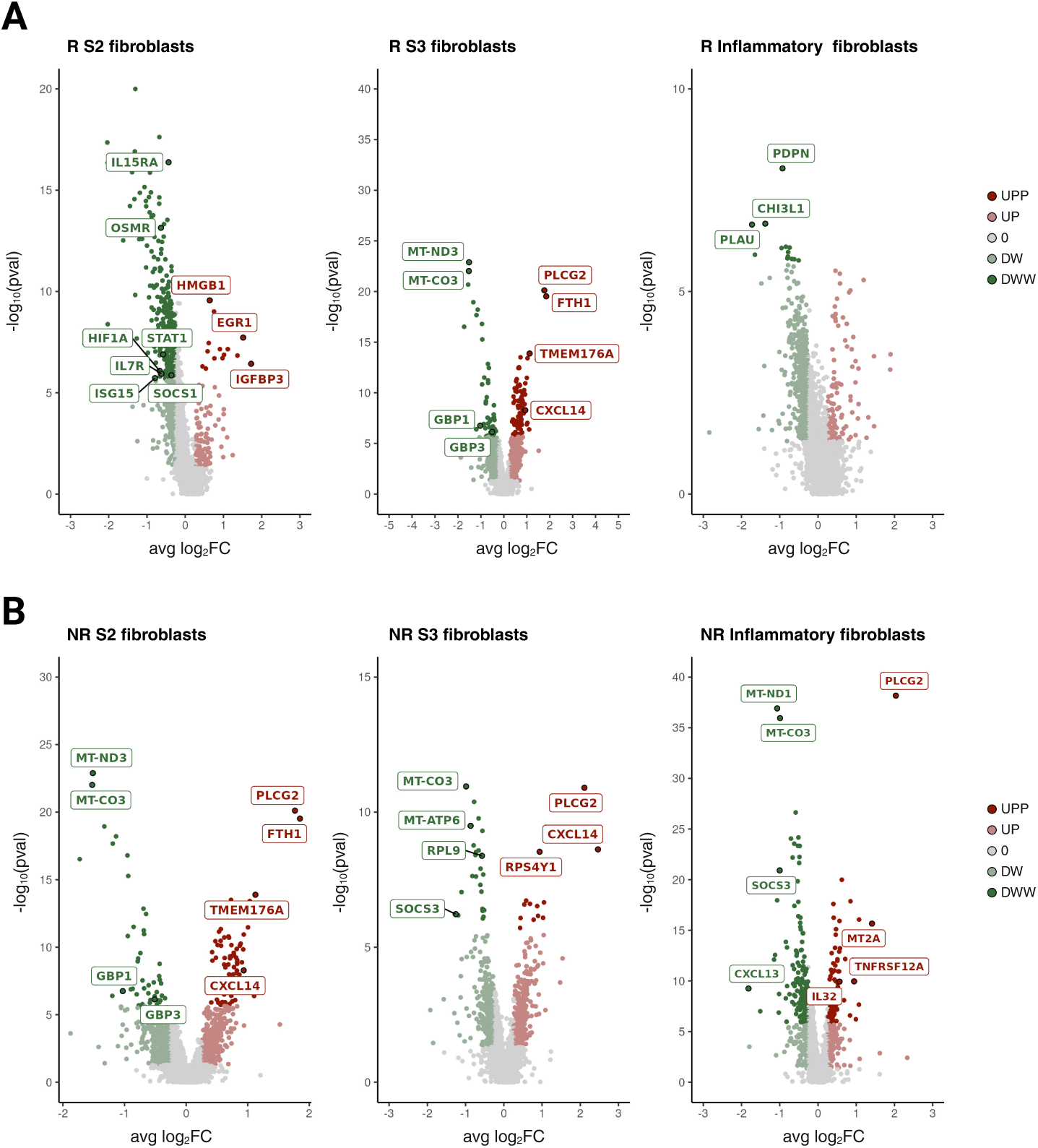
Tofacitinib induces transcriptional changes in subsets of intestinal fibroblasts. Volcano plots showing the main differentially expressed genes during treatment (post-tx *vs* pre-tx) in the macrophage subsets of responders (A) and non-responders (B). A two-sided Wilcoxon rank sum test was applied. Genes with a false discovery rate adjusted p value < 0.05, and a fold change (FC) > 1.2 (UPP, dark red) or FC < 0.83 (DWW, dark green) were considered regulated. Additionally, genes with a nominal p-value < 0.05 are shown in red (UP) if FC > 1.2 and green if (DW) FC < 0.83.

**Supplementary Figure 7.**
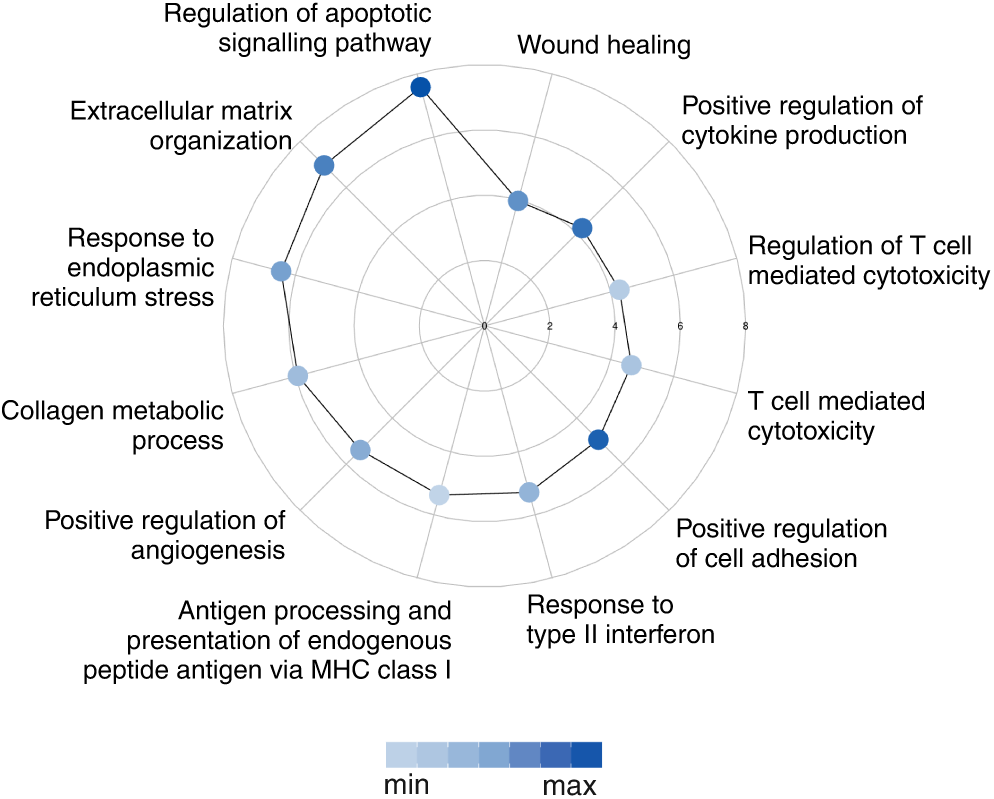
Tofacitinib regulates transcriptional pathways in S1 fibroblasts of responder patients. Polar plot showing significantly upregulated pathways in S1 fibroblasts from responders following treatment with tofacitinib. The radius of the plot represents - log10(p.value), while the color of the points indicates the gene ratio.

**Supplementary Figure 8.**
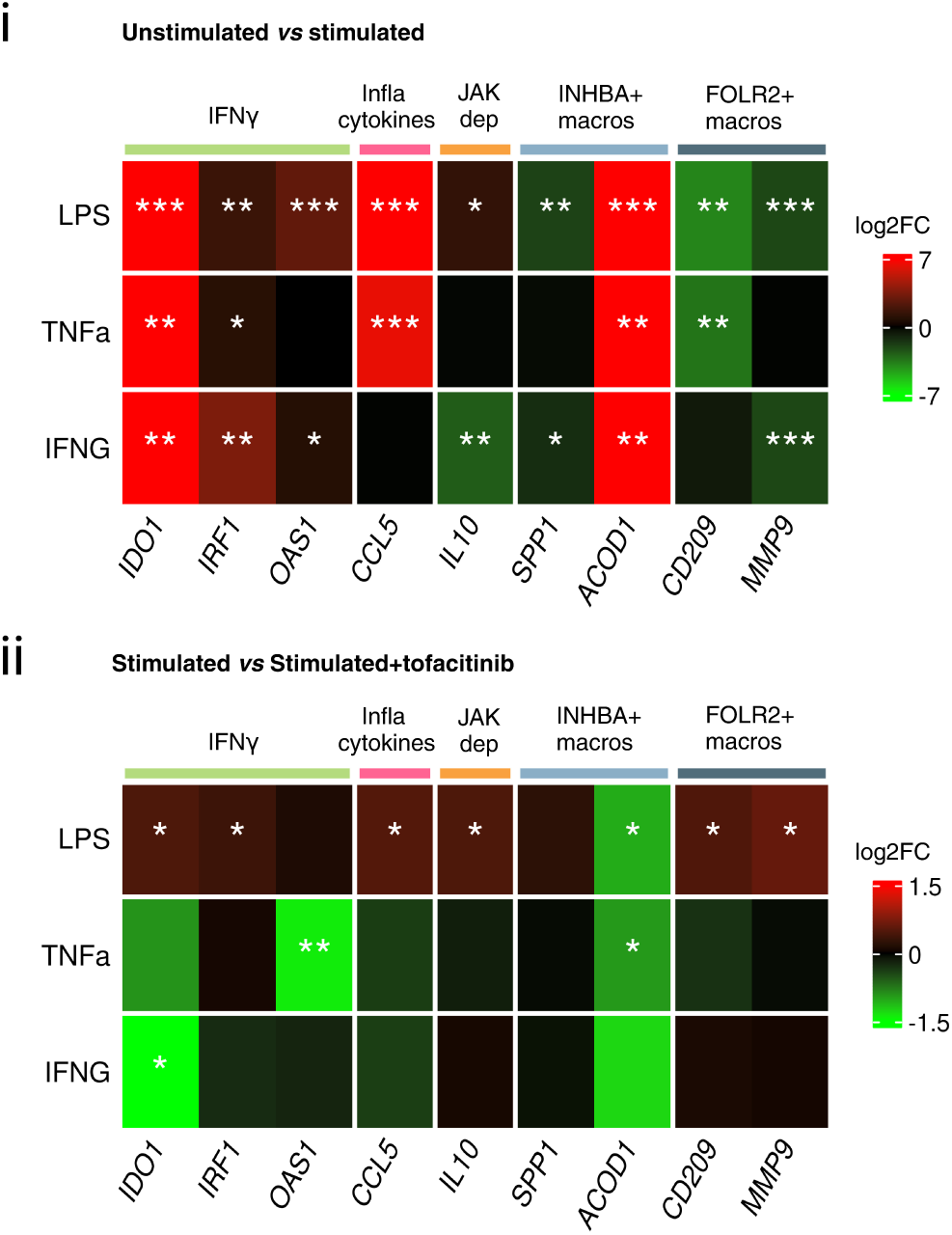
Effects of tofacitinib on macrophage activation. Heatmaps showing mean log2 of the fold-change (log2FC) in the expression of selected genes on monocyte-derived macrophages. i) Changes in response to LPS (10 ng/ml), TNF (20 ng/ml) and IFNγ (5 ng/ml) compared to unstimulated control are shown. ii) Changes in gene expression induced by tofacitinib (300 nM) on LPS, TNF or IFNγ−stimulated macrophages. infla, inflammatory; dep, dependent; macros, macrophages. n= 5 independent experiments. One-sample t-test false discovery rate-corrected p-values: *<0.05, **<0.01, ***<0.001, ****<0.0001.

**Supplementary Figure 9.**
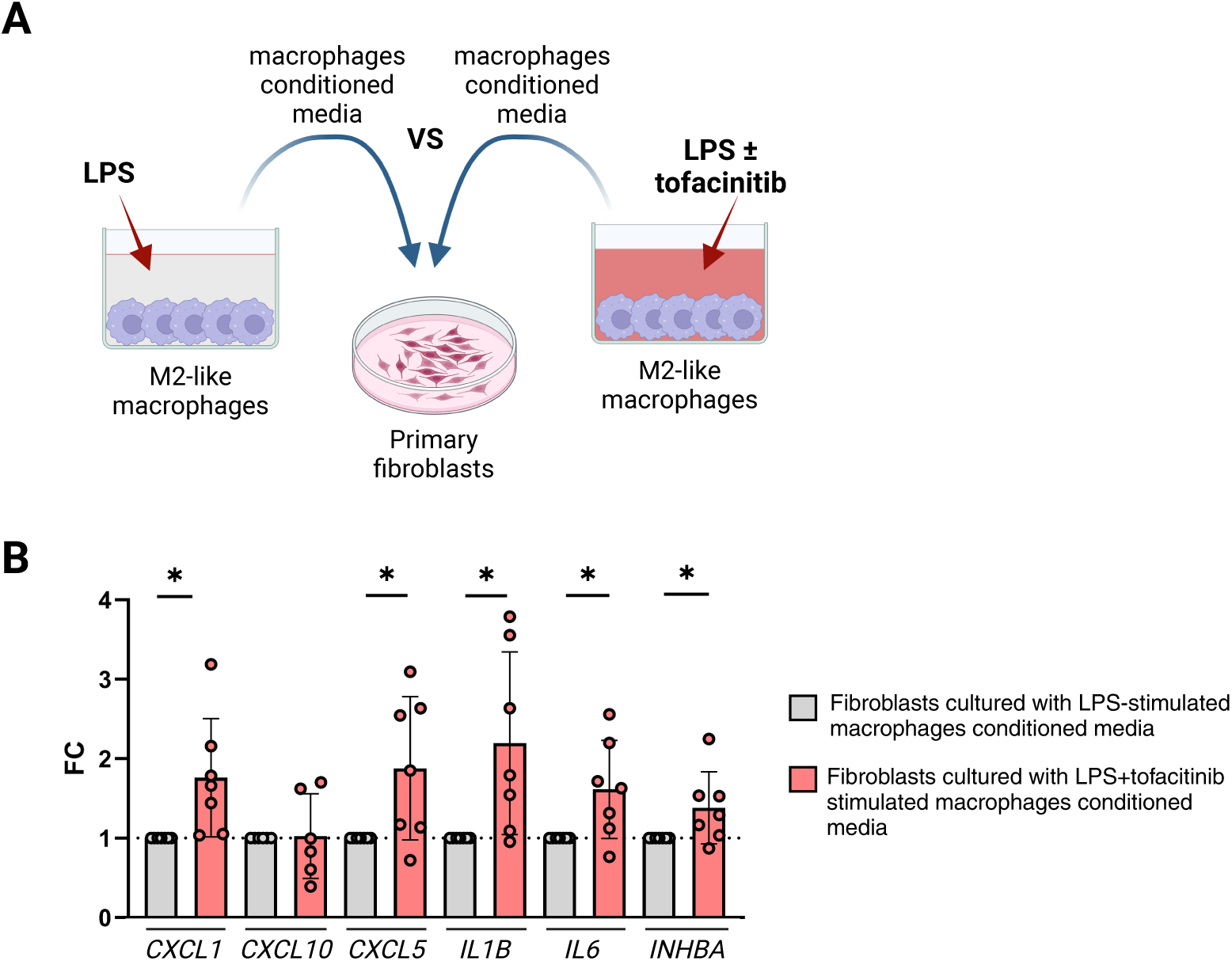
Effects of macrophage supernatants on intestinal fibroblasts. A) Schematic representation of experiment workflow. B) FC of mRNA gene expression of selected markers in colonic-derived fibroblasts incubated for 19h with conditioned media of LPS-stimulated macrophages in the presence of tofacitinib compared to conditioned media of LPS-stimulated macrophages treated with vehicle (DMSO). n=7 independent experiments. One-sample t-test false discovery rate-corrected p-1

## REFERENCES

1. Tsai L, et al. Contemporary Risk of Surgery in Patients With Ulcerative Colitis and Crohn’s Disease: A Meta-Analysis of Population-Based Cohorts. Clinical gastroenterology and hepatology : the official clinical practice journal of the American Gastroenterological Association. 2021;19(10):2031–45.

2. Tse CS, et al. Inflammatory Bowel Diseases-related Disability: Risk Factors, Outcomes, and Interventions. Inflamm Bowel Dis. 2023;30(3):501–7.

3. Sandborn WJ, et al. Tofacitinib as Induction and Maintenance Therapy for Ulcerative Colitis. N Engl J Med. 2017;376(18):1723–36.

4. Salas A, et al. JAK-STAT pathway targeting for the treatment of inflammatory bowel disease. Nat Rev Gastroenterol Hepatol. 2020;17(6):323–37.

5. Garrido-Trigo A, and Salas A. Molecular structure and function of Janus kinases: implications for the development of inhibitors. Journal of Crohn’s & colitis. 2019;14(Supplement_2):S713–S24.

6. Meyer DM, et al. Anti-inflammatory activity and neutrophil reductions mediated by the JAK1/JAK3 inhibitor, CP-690,550, in rat adjuvant-induced arthritis. J Inflamm (Lond). 2010;7:41.

7. Kapizioni C, et al. Biologic therapy for inflammatory bowel disease: Real-world comparative effectiveness and impact of drug sequencing in 13,222 patients within the UK IBD BioResource. Journal of Crohn’s & colitis. 2023;18(6):790–800.

8. Schubert M, et al. Perturbation-response genes reveal signaling footprints in cancer gene expression. Nat Commun. 2018;9(1):20.

9. Moore KW, et al. Interleukin-10 and the interleukin-10 receptor. Annu Rev Immunol. 2001;19:683–765.

10. Carl VS, et al. Role of endogenous IL-10 in LPS-induced STAT3 activation and IL-1 receptor antagonist gene expression. J Leukoc Biol. 2004;76(3):735–42.

11. Cuevas VD, et al. The Gene Signature of Activated M-CSF-Primed Human Monocyte-Derived Macrophages Is IL-10-Dependent. J Innate Immun. 2022;14(3):243–56.

12. Aschenbrenner D, et al. Deconvolution of monocyte responses in inflammatory bowel disease reveals an IL-1 cytokine network that regulates IL-23 in genetic and acquired IL-10 resistance. Gut. 2021;70(6):1023–36.

13. Alexander AF, et al. Single-cell secretion analysis reveals a dual role for IL-10 in restraining and resolving the TLR4-induced inflammatory response. Cell reports. 2021;36(12):109728.

14. Efremova M, et al. CellPhoneDB: inferring cell-cell communication from combined expression of multi-subunit ligand-receptor complexes. Nature protocols. 2020;15(4):1484–506.

15. Jiang P, et al. Systematic investigation of cytokine signaling activity at the tissue and single-cell levels. Nature methods. 2021;18(10):1181–91.

16. Singh S, et al. First- and Second-Line Pharmacotherapies for Patients With Moderate to Severely Active Ulcerative Colitis: An Updated Network Meta-Analysis. Clinical gastroenterology and hepatology : the official clinical practice journal of the American Gastroenterological Association. 2020;18(10):2179–91.

17. Planell N, et al. Transcriptional analysis of the intestinal mucosa of patients with ulcerative colitis in remission reveals lasting epithelial cell alterations. Gut. 2013;62(7):967–76.

18. Haberman Y, et al. Ulcerative colitis mucosal transcriptomes reveal mitochondriopathy and personalized mechanisms underlying disease severity and treatment response. Nat Commun. 2019;10(1):38.

19. Argmann C, et al. Biopsy and blood-based molecular biomarker of inflammation in IBD. Gut. 2023;72(7):1271–87.

20. Argmann C, et al. Molecular Characterization of Limited Ulcerative Colitis Reveals Novel Biology and Predictors of Disease Extension. Gastroenterology. 2021;161(6):1953–68

21. Arijs I, et al. Mucosal gene signatures to predict response to infliximab in patients with ulcerative colitis. Gut. 2009;58(12):1612–9.

22. Arijs I, et al. Mucosal gene expression of antimicrobial peptides in inflammatory bowel disease before and after first infliximab treatment. PloS one. 2009;4(11):e7984.

23. Gaujoux R, et al. Cell-centred meta-analysis reveals baseline predictors of anti-TNFalpha non-response in biopsy and blood of patients with IBD. Gut. 2018.

24. West NR, et al. Oncostatin M drives intestinal inflammation and predicts response to tumor necrosis factor-neutralizing therapy in patients with inflammatory bowel disease. Nat Med. 2017;23(5):579–89.

25. Friedrich M, et al. IL-1-driven stromal-neutrophil interactions define a subset of patients with inflammatory bowel disease that does not respond to therapies. Nat Med. 2021;27(11):1970–81.

26. Telesco SE, et al. Gene Expression Signature for Prediction of Golimumab Response in a Phase 2a Open-Label Trial of Patients With Ulcerative Colitis. Gastroenterology. 2018;155(4):1008–11.

27. Smillie CS, et al. Intra- and Inter-cellular Rewiring of the Human Colon during Ulcerative Colitis. Cell. 2019;178(3):714–30 e22.

28. Parikh K, et al. Colonic epithelial cell diversity in health and inflammatory bowel disease. Nature. 2019;567(7746):49-55.

29. Kinchen J, et al. Structural Remodeling of the Human Colonic Mesenchyme in Inflammatory Bowel Disease. Cell. 2018;175(2):372–86.

30. Kong L, et al. The landscape of immune dysregulation in Crohn’s disease revealed through single-cell transcriptomic profiling in the ileum and colon. Immunity. 2023;56(2):444–58.

31. Corridoni D, et al. Single-cell atlas of colonic CD8(+) T cells in ulcerative colitis. Nat Med. 2020;26(9):1480–90.

32. Boland BS, et al. Heterogeneity and clonal relationships of adaptive immune cells in ulcerative colitis revealed by single-cell analyses. Sci Immunol. 2020;5(50).

33. Huang B, et al. Mucosal Profiling of Pediatric-Onset Colitis and IBD Reveals Common Pathogenics and Therapeutic Pathways. Cell. 2019;179(5):1160–76.

34. Elmentaite R, et al. Single-Cell Sequencing of Developing Human Gut Reveals Transcriptional Links to Childhood Crohn’s Disease. Developmental cell. 2020;55(6):771–83.

35. Kanke M, et al. Single-Cell Analysis Reveals Unexpected Cellular Changes and Transposon Expression Signatures in the Colonic Epithelium of Treatment-Naïve Adult Crohn’s Disease Patients. Cell Mol Gastroenterol Hepatol. 2022;13(6):1717–40.

36. Ip WKE, et al. Anti-inflammatory effect of IL-10 mediated by metabolic reprogramming of macrophages. Science. 2017;356(6337):513–9.

37. Zigmond E, et al. Macrophage-restricted interleukin-10 receptor deficiency, but not IL-10 deficiency, causes severe spontaneous colitis. Immunity. 2014;40(5):720–33.

38. Shouval DS, et al. Interleukin-10 receptor signaling in innate immune cells regulates mucosal immune tolerance and anti-inflammatory macrophage function. Immunity. 2014;40(5):706–19.

39. Koelink PJ, et al. Anti-TNF therapy in IBD exerts its therapeutic effect through macrophage IL-10 signalling. Gut. 2020;69(6):1053–63.

40. Ghoreschi K, et al. Modulation of innate and adaptive immune responses by tofacitinib (CP-690,550). J Immunol. 2011;186(7):4234–43.

41. Thomas T, et al. A longitudinal single-cell therapeutic atlas of anti-tumour necrosis factor treatment in inflammatory bowel disease. bioRxiv. 2023;10.1101/2023.05.05.539635.

42. Khanna D, et al. Tofacitinib blocks IFN-regulated biomarker genes in skin fibroblasts and keratinocytes in a systemic sclerosis trial. JCI Insight. 2022;7(17):e159566.

43. Garrido-Trigo A, et al. Macrophage and neutrophil heterogeneity at single-cell spatial resolution in human inflammatory bowel disease. Nat Commun. 2023;14(1):4506.

44. Mereu E, et al. Benchmarking single-cell RNA-sequencing protocols for cell atlas projects. Nat Biotechnol. 2020;38(6):747–55.

45. Love MI, et al. Moderated estimation of fold change and dispersion for RNA-seq data with DESeq2. Genome Biol. 2014;15(12):550.

46. Butler A, et al. Integrating single-cell transcriptomic data across different conditions, technologies, and species. Nat Biotechnol. 2018;36(5):411–20.

47. Yu G, et al. clusterProfiler: an R package for comparing biological themes among gene clusters. OMICS. 2012;16(5):284–7.

48. Puig-Kroger A, et al. Folate receptor beta is expressed by tumor-associated macrophages and constitutes a marker for M2 anti-inflammatory/regulatory macrophages. Cancer Res. 2009;69(24):9395–403.

